# How accurate is genomic prediction across wild populations?

**DOI:** 10.1101/2025.06.16.659953

**Authors:** Kenneth Aase, Hamish A. Burnett, Henrik Jensen, Stefanie Muff

**Affiliations:** Department of Mathematical Sciences, Norwegian University of Science and Technology, Trondheim, Norway; The Gjærevoll Centre, Norwegian University of Science and Technology, Trondheim, Norway; Department of Biology, Norwegian University of Science and Technology, Trondheim, Norway

**Keywords:** accuracy, across-population, genomic prediction, house sparrow, population differentiation, wild populations

## Abstract

Evolutionary ecology seeks to understand causes and consequences of evolutionary changes across time and space, and genomic data present novel opportunities to investigate these processes. Genomic prediction – that is, predicting individual genetic values from high-density marker data – has revolutionized breeding programs and medical genetics. In wild populations, however, genomic prediction has been used in only a handful of studies, and only *within* populations. There is still a lack of applications that predict across wild populations, which could provide answers to questions related to spatially varying evolutionary processes, such as local adaptation. A severe challenge for across-population genomic prediction, however, is the decrease in accuracy when models are trained on data from one population and predict genetic values in another. Here, we used genomic prediction to predict across wild house sparrow populations, and compared the accuracy to within-population prediction. Predictions across populations were less accurate and more variable than within populations. We also highlighted limitations of the current theory for general genomic prediction accuracy, and related across-population accuracy to several population differentiation measures. Our results underline the necessity of understanding why genomic prediction currently performs poorly across populations, and of developing methods that exploit genomic data in novel ways.

## Introduction

Quantifying the genetic basis of a given phenotypic trait is useful for understanding how evolutionary processes act in nature (Lynch & Walsh, 1998; Kruuk *et al*., 2008; Charmantier *et al*., 2014; Walsh & Lynch, 2018). In wild populations, the prediction of an individual’s genetic value (also known as breeding value), which is the additive contribution of the individual’s genome to its phenotype (Hill, 2014), can help us track adaptive and non-adaptive evolutionary changes across time, detect local adaptation to spatially varying environmental conditions, investigate selection processes, make inferences about unmeasured or unexpressed phenotypes in genotyped individuals, and compare how much individuals’ genotypes contribute to trait variation relative to environmental conditions (Jensen *et al*., 2014; Hunter et al., 2022). Traditionally, genetic values for complex traits have been predicted using relatedness information from multi-generational pedigrees (Henderson, 1984; Kruuk, 2004; Kruuk *et al*., 2008), but thanks to advances in genotyping technologies we are now less reliant on the labor- and time-consuming task of constructing pedigrees from long-term study systems. Methods that instead rely on genomic data, usually individual genotype information on high-density genome-wide sets of single-nucleotide polymorphism (SNP) markers, to predict genetic values are known as *genomic prediction* (GP).

The idea of GP originated in animal breeding (Meuwissen *et al*., 2001), where the goal is artificial selection on profitable traits. By exploiting genomic data from the animals, the genetic values can be predicted already in early life stages, speeding up the selection process by eliminating the need to wait for a trait to be expressed in the individual itself or its descendants. Additionally, GP is expected to more accurately predict genetic values compared to pedigree-based methods (Gienapp *et al*., 2017).

Due to these factors, and the decreasing cost of SNP genotyping, GP is now widely used in animal and plant breeding (Meuwissen *et al*., 2016; Hickey *et al*., 2017; Crossa *et al*., 2017). Furthermore, medical geneticists are developing *polygenic risk scoring*, a variation of GP that shows promise for use in personalized medicine to detect an individual’s genetic risk for complex diseases (Khera *et al*., 2018; Wray *et al*., 2019; Choi et al., 2020).

In wild populations, application of GP is still in its infancy (McGaugh *et al*., 2021), possibly due to additional challenges present in wild study systems compared to breeding or medical genetics. Firstly, high-density genomic data sets are less common and sample sizes are generally smaller than in the aforementioned fields, because of inherent difficulties and higher costs of sampling wild animals. Second, the *effective* population sizes *N*_*e*_ are often larger in wild systems than in agriculture, making it more challenging to obtain accurate and precise estimates of additive genetic effects (Gienapp *et al*., 2019). A third important challenge is the need to account for environmental heterogeneity and population structure (Kruuk & Hadfield, 2007; Aase *et al*., 2022), both of which are common in wild populations, but outside the researcher’s control. Given these challenges, it is not surprising that only a few GP studies have been performed in wild animal populations, for example, in great tits (*Parus major*, see Gienapp *et al*., 2019; Verhagen *et al*., 2019; Lindner *et al*., 2023, 2024), Soay sheep (*Ovis aries*, see Ashraf *et al*., 2022; Hunter et al., 2022; Vahedi et al., 2023), red deer (*Cervus elaphus*, see Gauzere *et al*., 2023), three-spine stickleback (*Gasterosteus aculeatus*, see Strickland *et al*., 2024) and house sparrows (*Passer domesticus*, see Aspheim *et al*., 2024). A shared methodological characteristic among these studies was that they all used GP to predict the genetic values of individuals sampled from the same (meta-)population as the individuals used to train (*i*.*e*., fit) the model. However, many potential applications of GP are reliant on prediction from one population into *another* population.

In light of the high cost and aforementioned difficulty of sampling wild populations, borrowing information from phenotyped populations to make inferences about non-phenotyped individuals in other populations would be a very useful application of GP. *Across-population* GP here refers to models and applications in which the phenotyped and genotyped individuals used to fit the model and the unmeasured, but genotyped, individuals for which we predict genetic values are sampled from two different populations. With reliable across-population GP methods one would be able to train a GP model for hard-to-measure phenotypes in one population, and only need to genotype another population to make inferences about this population’s genetic values, greatly expanding the number of populations where we are able to answer the aforementioned evolutionary and quantitative genetic questions. Across-population GP could also be especially relevant in conservation programs that aim to use captive breeding and translocation to strengthen existing natural populations (*e*.*g*., Sauve *et al*., 2022), where the predicted fitness of an individual in another population is of interest. Similarly, GP may also be useful across temporal, as opposed to spatial, distances between the training and test samples (Habier *et al*., 2007). Contemporary genotype and phenotype data coupled with ancient genetic material could allow for inferences about ancestors that are temporally distant and provide information on the rate and direction of any evolutionary change that has occurred within the population over time (*e*.*g*., Berens *et al*., 2017; Cox et al., 2019; Marciniak et al., 2022).

A major problem, known from both agricultural and medical applications, is that GP across populations (*i*.*e*., breeds or ethnicities) suffers from seriously degraded prediction accuracy compared to within-population GP (Habier *et al*., 2007; Hayes *et al*., 2009a; Martin *et al*., 2017, 2019; Rio *et al*., 2021; Lupi et al., 2024). Similar issues have been discussed in the context of using polygenic scores in paleogenetics, where the target population is ancient, and thus very temporally distant from contemporary training data (Irving-Pease *et al*., 2021; Carlson *et al*., 2022). Possible explanations for this phenomenon include population-differences in allele frequencies or linkage disequilibrium (LD) patterns, lack of family relationships, or differences in allele effects caused by non-additive genetic effects and/or genotype-by-environment interactions (Zhong *et al*., 2009; Wang et al., 2020; Hou et al., 2023).

More generally, a theoretical understanding of expected GP accuracy in a broad range of scenarios would be essential. Some previous work was targeted at theoretically understanding and forecasting the GP accuracy in given scenarios (Daetwyler *et al*., 2008; Goddard et al., 2011; Lee et al., 2017; Dekkers *et al*., 2021), and how much the accuracy is expected to degrade when predicting across populations (Wientjes *et al*., 2015; Wang *et al*., 2020; Ding *et al*., 2023). When planning studies, for example, such deterministic formulas can help decide on the required amount of field sampling and genotyping, and can allow one to assess *a priori* whether a planned experiment or study has the power to deliver the desired insight. However, as we will discuss, formulas for expected accuracy suffer from various deficiencies, especially when applied across populations, and so the problem of across-population GP accuracy still needs further investigation.

Wild systems differ from the populations used in animal breeding and human genetics in many ways. Thus, how well across-population GP works in wild populations has not previously been examined, and the challenges with across-population GP are not well known within evolutionary ecology. In order to improve our understanding of how well standard methods for GP work when used across populations, we here compared the accuracy of within- and across-population GP for morphological traits of wild house sparrow populations. The island structure of the study populations, along with the relatively large sample sizes from this long-term study system, makes this data set uniquely suited to explore any challenges of across-population GP. We compared the performance of GP within and across populations, and explored whether various measures of genetic differentiation between populations affected the across-population accuracy. Additionally, we discussed various aspects of GP accuracy, including proper scaling methods for accuracy and how to deal with repeated measurements, and we demonstrated that existing formulas for expected accuracy are inappropriate for across-population GP.

## Methods and materials

### House sparrow data

The data underlying all GP analyses presented here stem from long-term individual-based house sparrow study populations located on several islands along the coast of Norway (see Figure 1 and Jensen *et al*., 2013), including 12 islands in an insular meta-population on the coast of Helgeland (Sæther *et al*., 1999; Ranke et al., 2021) and 4 islands 75-295 km south of this meta-population (Kvalnes *et al*., 2017; Ranke et al., 2020; Nafstad et al., 2023). The long-term nature of the study (most populations have been studied since 1993), along with high capture rates, relatively low levels of natal dispersal and virtually no breeding dispersal (Ranke *et al*., 2021), makes it attractive for investigating evolutionary ecological hypotheses. When the sparrows are first captured, they are marked with a numbered metal ring and three colored plastic rings, which uniquely identifies them in future recaptures and re-sightings. Birds first ringed as nestlings or fledged juveniles in the summer were known to have hatched that year. Because of the high ringing and recapture rate in our study populations birds first captured in the autumn were assumed to have hatched the same year as they were first captured, and birds first captured as adults from January to August were assumed to have hatched the previous year (Niskanen *et al*., 2020; Araya-Ajoy et al., 2021). The study populations have been subject to an extensive pheno- and genotyping effort, with a total of over 12 000 house sparrows genotyped on either a custom Axiom 200K SNP array (Lundregan *et al*., 2018; Niskanen et al., 2020) or a cross-compatible Axiom 70K SNP array containing a subset of the SNPs on the 200K Array with identical probe design (Burnett *et al*., in prep.). A subset of these genotyped individuals were recorded as adults and measured for various morphological traits, including body mass (to the nearest 0.1 gram), tarsus length (to the nearest 0.01 millimeter) and wing length (to the nearest millimeter), with repeated measurements over several years available for many of the individuals and measurements adjusted for any fieldworker differences (see *e*.*g*. Niskanen *et al*., 2020).

**Figure 1:**
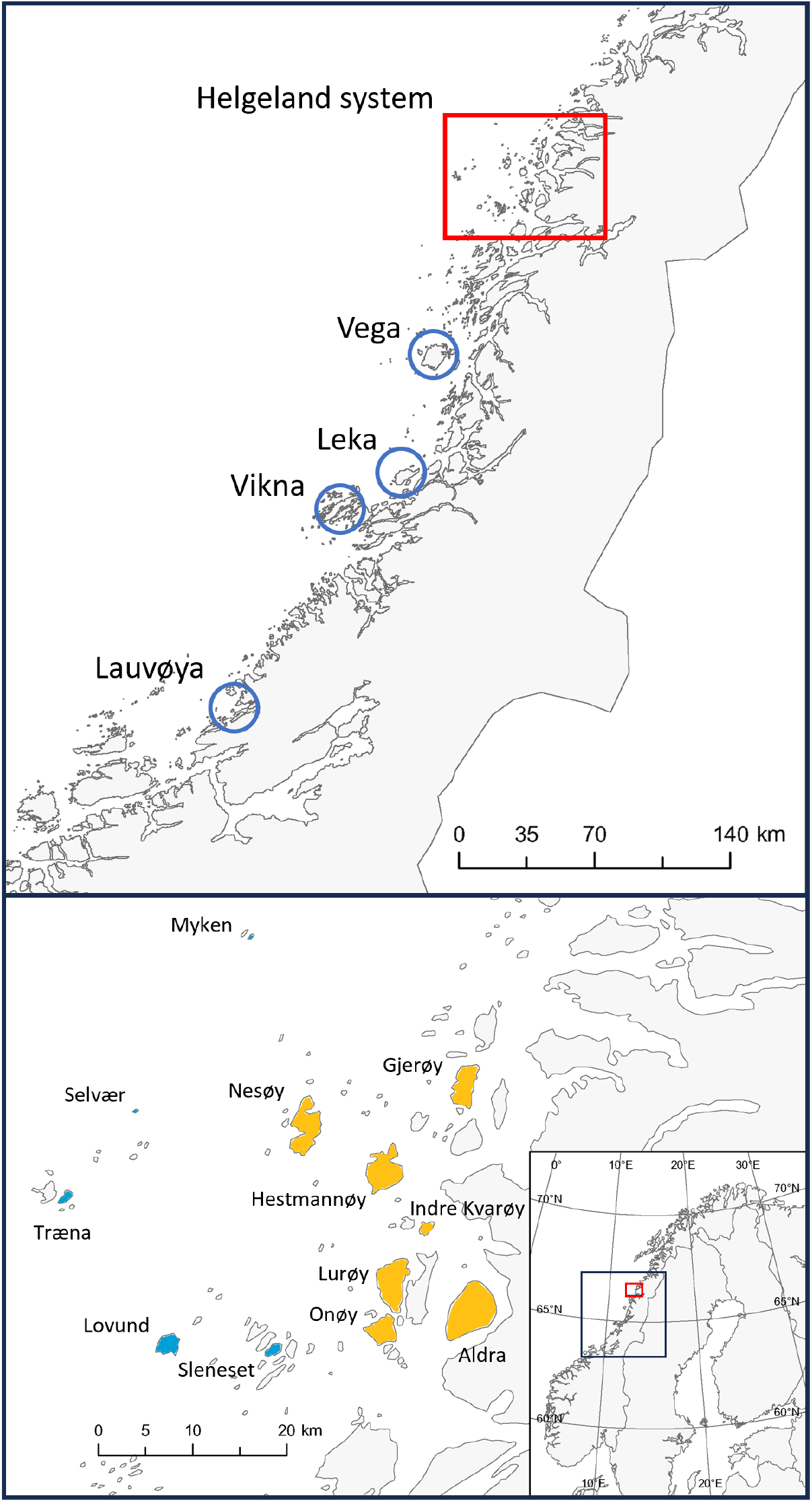
Maps showing the locations of the various island populations in the study. In the top panel, the Helgeland system is indicated with a red square, while the blue circles indicate the southern islands of Vega, Leka, Vikna and Lauvøya. The bottom panel shows the Helgeland system, with farm islands colored yellow and non-farm islands colored blue. The map in the bottom right shows the location of the other two maps, zoomed out to the scale of Norway.

The hierarchical geographic island structure in the study systems makes the house sparrow data ideal for investigating the accuracy of across-population GP. We leverage this natural structure to investigate a range of situations relevant for wild populations in general. As explained below, we organize the data into three scenarios with increasingly stronger population structure (Jensen *et al*., 2013; Nafstad *et al*., 2023; Ranke et al., 2024): the Helgeland meta-population (*Scenario 1*, islands connected through dispersal), the southern islands (*Scenario 2*, islands connected through translocations), and a merged scenario (*Scenario 3*) where we use all the available data from Scenarios 1 and 2. Overall sample sizes for each/trait scenario are shown in Table 1, though each GP model was trained on a subset of these samples, as explained below.

**Table 1:** The number of unique individuals that were genotyped and measured for a given phenotype. The numbers in the parentheses are the total number of measurements, which is higher than the number of unique individuals due to repeated measurements.

| Trait | Scenario |  |  |
| --- | --- | --- | --- |
|  | 1: Helgeland | 2: Southern | 3: Merged |
| Body mass | 3456 (7977) | 2254 (4124) | 5710 (12101) |
| Tarsus length | 3445 (8067) | 2255 (4171) | 5700 (12238) |
| Wing length | 3424 (7997) | 2254 (4151) | 5678 (12148) |

### Scenario 1: Helgeland

The first scenario is based on the data from the Helgeland system, a meta-population located in an archipelago off the coast of Northern Norway. We used data collected on the islands in the archipelago which are named in the map in the lower panel in Figure 1. Although the house sparrow is generally a sedentary bird species, the island populations in the Helgeland archipelago are connected by the sparrows that disperse between the islands (Ranke *et al*., 2021; Saatoglu *et al*., 2021). The islands are thus genetically connected by spatially varying degrees of dispersal, resulting in varying levels of genetic differentiation between them. Notably, the islands can broadly be categorized into “farm” islands, where the sparrows usually nest in the barns of dairy farms, or “non-farm” islands, where the sparrows live in local peoples’ gardens where their environment is generally harsher outside the breeding season. Previous studies have found that individuals from these two island types are phenotypically and genetically differentiated (Holand *et al*., 2011; Jensen et al., 2013; Saatoglu et al., 2024), and in particular that they differ in both the means and variances of the genetic values for various morphological traits (Muff *et al*., 2019; Aase et al., 2022).

### Scenario 2: Southern

The second scenario corresponds to four “farm” islands located south of the Helgeland meta-population (Vega, Leka, Vikna and Lauvøya; see Figure 1, upper panel). Due to the distances between these four islands, there is virtually no natural dispersal between them (Ranke *et al*., 2024). However, some of the southern island populations have been connected by past translocation experiments, in which individuals were moved between the islands (for details, see Kvalnes *et al*., 2017; Ranke *et al*., 2020; Nafstad *et al*., 2023). In other words, Scenario 2 consists of separate, but partially connected island-populations, and thus has stronger population structure than the Helgeland scenario (Jensen *et al*., 2013).

### Scenario 3: Merged

In the third scenario, denoted *merged*, we used all the data from the two first scenarios (Helgeland system and southern islands) simultaneously. Estimates of genetic differentiation suggest low dispersal rates between the Helgeland system and the southern islands (Jensen *et al*., 2013), which is supported by only a handful of dispersal events recorded across more than 20 years by recapture and re-sighting of thousands of ringed house sparrows (Ranke *et al*., 2024). Thus, in practice, the southern islands are wholly separate populations from the Helgeland meta-population, so the data in Scenario 3 have a higher hierarchical level of structure than Scenarios 1 and 2.

## Statistical methods

### Models for genomic prediction

All GP models were formulated as mixed models with the aforementioned morphological traits (body mass, tarsus length, or wing length) as continuous responses, given as

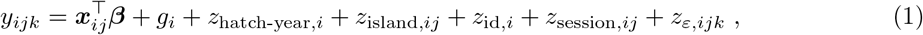

where *y*_*ijk*_ is the *k*^th^ phenotypic measurement of individual *i* during *measurement session j* (see below), ***x***_*ij*_ is a vector of fixed covariates with corresponding fixed effects vector ***β***, encoding for an intercept term, a binary variable for phenotypic sex (0 = female, 1 = male), a categorical variable for the month of measurement (from January = 1 through November = 11), and a quantitative variable for age in years (*i*.*e*., the difference between measurement year and hatch year; range 1-11). The other terms in Equation (1) are random effects, with *g*_*i*_ being a structured Gaussian random effect representing the genetic value of individual *i*, and the remaining terms being independent unstructured Gaussian random effects accounting for various confounding effects. The random effect *z*_hatch-year,*i*_ accounted for the effect of individual *i*’s hatch year (range 1992–2020), *z*_island,*ij*_ accounted for the effect of island of measurement (*i*.*e*., one of the islands named on Figure 1), and *z*_id,*i*_ accounted for remaining permanent environmental effects specific to individual *i*. Following Ponzi et al. (2018), we defined a measurement session to be the set of measurements of a given individual made on the same day, and included the measurement session-specific random effect *z*_session,*ij*_. Since the birds experience essentially the same environment during a measurement session, *z*_session,*ij*_ thus accounted for all residual variation except measurement error, which is captured by *z*_*ε,ijk*_. We fitted the mixed models in a Bayesian framework using R-INLA (Rue *et al*., 2009), and assigned independent N (0, 10^3^) priors to all entries in ***β***. For all random effect variances we assigned independent penalized complexity priors 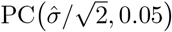, where 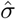 denotes the sample standard deviation of the respective model’s response in the training set (Simpson *et al*., 2017). In other words, we ascribed a prior probability of 5% that a given random effect would explain more than half the sample variance.

We modeled the genetic value *g*_*i*_ using a genomic animal model formulation, that is, we assumed that the vector 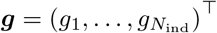 follows a normal distribution 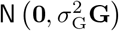, where *N*_ind_ is the number of individuals, 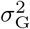 is the additive genetic variance (often denoted *V*_*A*_) and **G** is an *N*_ind_ *× N*_ind_ genomic relatedness matrix (GRM). To estimate the relatedness between individuals *i* and *i*′ (*i*.*e*., entry *G*_*ii*_*′* of **G**) we used the estimator

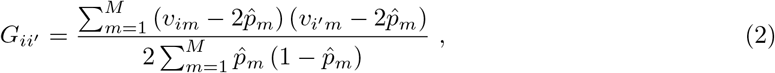

as given by VanRaden (2008), where *M* is the number of SNPs, *v*_*im*_ is the number (0, 1 or 2) of minor alleles individual *i* has at SNP *m* and 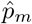 is the estimated minor allele frequency (MAF) at SNP *m*. Note that we always computed the allele frequencies 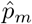 from the full population in the given scenario (Helgeland, southern or merged), which implies that the base population for estimates of 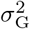 does not necessarily equal the ensemble of individuals in the combined training and test sets (Hayes *et al*., 2009b; Legarra, 2016). Furthermore, we ensured the positive-definiteness of **G** (Hollifield *et al*., 2022) by adding the lowest eigenvalue *d* of the given GRM to its diagonal (the exact values added ranged from 10^*−*9^ to 0.0066, see the tables in Appendix S6 for exact values in a given model).

Before we estimated the GRMs **G** as described above, we used `PLINK 1.9` (Chang *et al*., 2015) to apply quality control filters to the SNP data. Individuals were filtered for call rate (*>* 0.95), while SNPs were filtered for call rate (*>* 0.9) and MAF (*>* 0.01). The remaining missing variant calls were not imputed, but terms in Equation (2) where either *v*_*im*_ or *v*_*im*_*′* was a missing call were skipped in the computation of the entries of **G** (Chang, 2024, personal communication). The quality control filters were always applied twice: once for the scenario overall, and again for the subset of data used in a specific GP model (see below). Thus, the exact subset of SNPs and individuals differ slightly between different models (see the tables in Appendix S6 for the exact sample sizes and numbers of SNPs in each model).

### Within- and across-population genomic prediction

Generally in statistical prediction models, we distinguish between the data used to fit the model, which we label the *training set*, and the data used to assess the model, which we label the *test set*. Assessment of the prediction accuracy on the test set thus reflects the model’s performance on unseen data. How much information the training set provides about the test set is crucial in determining the performance of a prediction model, and the same is true in GP (Pszczola *et al*., 2012; Akdemir & Isidro y Sánchez, 2019; Rio et al., 2021). As the goal of this study is to demonstrate how across-population GP accuracy is affected by various factors, we draw extra attention to the design of our training and test sets. In each of the three scenarios we considered two types of models: *within*-population GP and *across*-population GP. In within-population GP, the training and test samples are drawn from the same (meta-)population, whereas in across-population prediction the training and test samples come from two distinct (meta-)populations (Figure 2).

**Figure 2:**
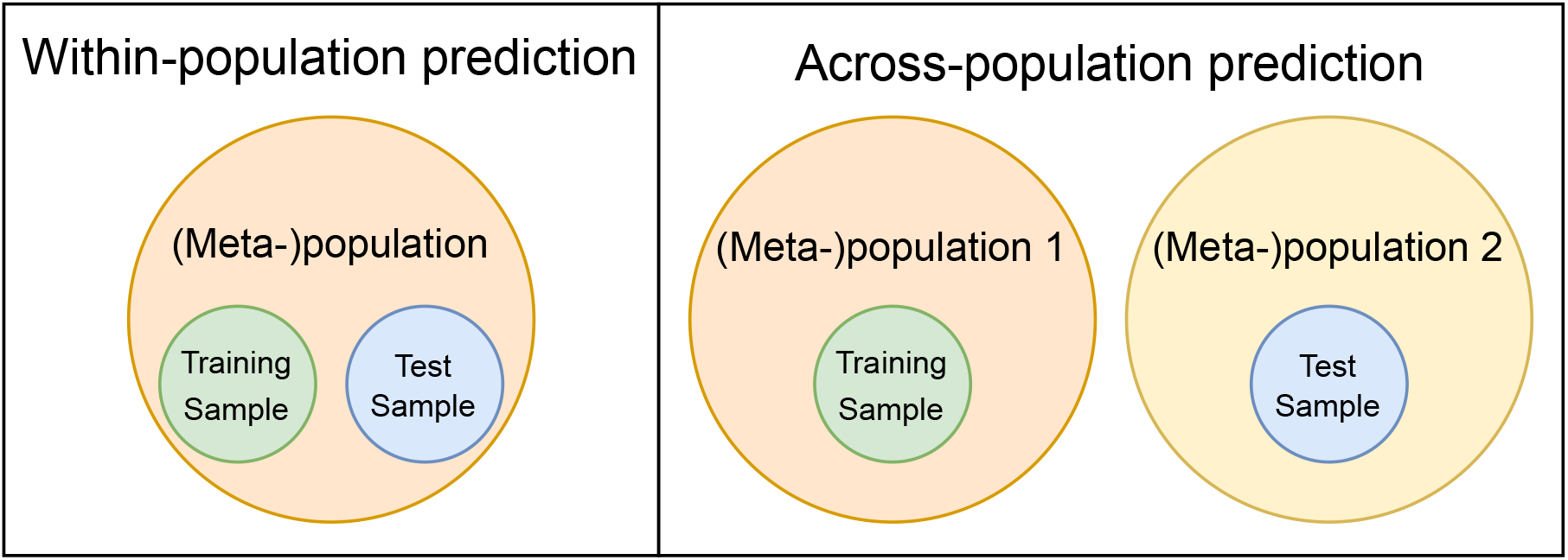
Conceptual diagram showing the difference between within-population and across-population GP. In within-population GP, the training and test samples are drawn from the same (meta-)population, whereas in across-population GP the training and test sample are drawn from two different (meta-)populations.

We assessed the within-population models using 10-fold cross-validation (CV), that is, by randomly partitioning the data in a given scenario into 10 equally-sized folds, and then fitting 10 GP models where in a given model one of the folds was used as the test set, and the remaining 9 were together used as the training set. Our within-population GP models thus emulated predicting the genetic values of genotyped but non-phenotyped individuals from an already-sampled (meta-)population. Aside from partitioning the folds by individuals rather by single measurements to prevent repeatedly measured individuals from ending up in more than one fold, we ignored the presence of any population structure when creating the folds for within-population GP. Thus, sparrows from all the different islands in a given scenario appeared in both the training and test set for a given within-population model. Notably, this approach causes closely related individuals to end up in different folds, giving high relatedness between training and test set individuals, and ensures that the environments experienced by the test set individuals were also represented in the training set.

Conversely, for our across-population models we drew the training and test individuals from different (meta-)populations. In particular, in Scenario 1 (Helgeland) we performed a leave-island-out CV, where for a given model all sparrows from one island made up the test set, and the sparrows from all other islands were used to train the model. We also fitted models in Scenario 1 where the farm islands made up the test set and the non-farm islands made up the training set, and vice versa, as well as models which predicted the genetic values for individuals from a given test island based on the data from all the farm islands (except the test island if it was a farm island), and based on the data from all the non-farm islands (except the test island if it was a non-farm island). For Scenario 2 (southern islands), we again performed a leave-island-out CV. In Scenario 3 (merged) we predicted the genetic values in the full Helgeland meta-population based on all the sparrows from the southern islands, and vice versa. Thus, the across-population models in the three scenarios captured varying degrees of differentiation between the (meta-)populations across which we predicted, both environmentally and genetically. Note that most populations contained some level of admixture (Aase *et al*., 2022), due to either natural dispersal or translocation (Kvalnes *et al*., 2017; Ranke et al., 2020, 2021; Saatoglu et al., 2021; Nafstad et al., 2023). There was thus some ambiguity about whether to assign individuals that were recorded on multiple islands to the test or training set in a given model. We dealt with this issue by using individuals’ hatch island to partition the full set of individuals into disjoint sub-populations, and letting the test set of a given model be the set of sparrows with hatch island corresponding to the test set islands. For sparrows that were observed as hatchlings, the hatch island is known. Otherwise we used the first island of observation as the sparrow’s hatch island, as this likely to be the individual’s true hatch island due to the sedentary nature and relatively low dispersal rate in house sparrows (Ranke *et al*., 2021; Saatoglu et al., 2021).

### Accuracy metric

Fitting model (1) in a Bayesian framework provided estimates of the full posterior distributions for all genetic values *g*_*i*_, and we used the posterior means, denoted *ĝ*_*i*_, as point estimates of *g*_*i*_. A common accuracy measure for GP models is

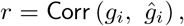

that is, the correlation between true and estimated genetic values in the test set of a given model. However, in real data studies the true *g*_*i*_ is an unknown latent variable and is not explicitly measurable. Therefore, one cannot *directly* compute *r*, except in simulation studies where *g*_*i*_ would be known.

Fortunately, scaling the easily estimable correlation Corr(*y*_*ijk*_, *ĝ*_*i*_) by an appropriate factor allows us to indirectly estimate *r*, given that the model is correctly specified (see Appendix S2). In the literature the scaling factor is commonly equated to *h*, the square root of narrow-sense heritability of the focal trait. However, as we show in Appendix S2, the appropriate definition of the factor should instead be

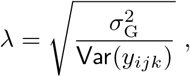

which does not necessarily equal *h* (since only biologically relevant sources of variation should be included in the denominator of *h*^2^, see Wilson, 2008; de Villemereuil *et al*., 2018, whereas *λ* also contains measurement error). We therefore measured the GP accuracy *r* in a given test set as

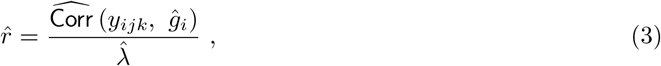

where 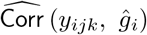 denotes the sample correlation between phenotypic measurements and estimated genetic values in the test set, and 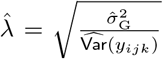, with 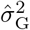 and 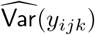 being estimates of 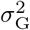 and the variance of the phenotypic measurements, respectively. To estimate 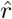 as accurately as possible, both 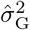 and 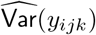 correspond only to individuals in the test set for a model. For the 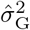 this was achieved by fitting separate genomic animal models with the full data set of a given scenario, and then subsampling the posteriors for individuals in the test set (see Appendix S1), while 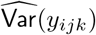 is the sample variance of measurements in the test set.

Note that the two vectors containing the realizations of the random variables *y*_*ijk*_ and *ĝ*_*i*_, that is, the phenotypic measurements and point estimates for *g*_*i*_, respectively, have different lengths due to the repeated measurements of the response. Therefore, when we estimated the sample correlation 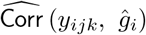, we increased the length of the vector of realizations of *ĝ*_*i*_ by appropriately repeating elements to match the length of the vector of realizations of *y*_*ijk*_, so that the indices *i* correspond. However, this repetition of elements causes 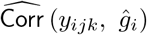 to be attenuated (biased towards zero, see *e*.*g*. Carroll *et al*., 1995). Fortunately, dividing by 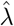 as suggested in equation (3) properly corrects for the attenuation in the correlation estimate (demonstrated in Appendix S2). Dividing by *λ* thus simultaneously provides an indirect estimator for *r*, and also corrects for the correlation-attenuation induced by repeated measures.

In addition to the point estimates 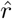 of the prediction accuracies, one can also calculate the uncertainty in 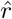. For the within-population models we can use the CV standard deviation, but the method for attaining the uncertainties in 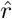 for the across-population models is more involved and involves resampling from the GP model posteriors. We describe this resampling-based method in Appendix S2.

## Formulas for expected accuracy

When planning or assessing a GP study, it may be useful to forecast how high the accuracy will be in a given scenario. Such theoretical *a priori* estimates of the GP accuracy can tell us how much data we need to collect to attain a certain level of accuracy, or whether the model is performing as well as expected. To this end, heuristic formulas that quantify E(*r*), the *expected* accuracy from a GP model, have been developed. Most such expressions are variations on the formula

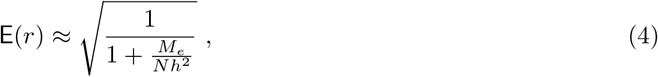

originally presented by Daetwyler et al. (2008, 2010), where *h*^2^ is the heritability of the trait, *N* is the number of individuals in the training set and *M*_*e*_ is the *effective number of independent chromosome segments*. The parameter *M*_*e*_ can thus be interpreted as the number of independent effects the model has to estimate from the data, and is positively related to effective population size and the length of the genome (Brard & Ricard, 2015). Alternatively, *M*_*e*_ is sometimes instead viewed as a model-specific parameter capturing how suitable the training set is for a given test set (Lee *et al*., 2017).

At first glance, formulas like Equation (4) appear to be helpful tools in assessing our GP model accuracies (as has been done before in wild systems, see Ashraf *et al*., 2022). However, use of formulas for expected accuracy is in practice severely complicated by various interconnected issues that we outline below. One set of issues relates to the underlying assumptions of the formula, while another involves the interpretations of the formula parameters. Additional issues arise in across-population GP. As we will argue, these issues necessitate that the user makes some arbitrary choices, which can potentially result in Equation (4) failing to predict the realized GP accuracies (as demonstrated in Appendix S3). What follows is an extended discussion of why we therefore prefer not to compare our observed accuracies (3) to the expected accuracies from (4).

### Issues related to formula assumptions

Equation (4), and its aforementioned generalizations, impose strict assumptions of an “ideal” scenario, namely an unstructured population of unrelated individuals sampled once in a homogeneous environment (Daetwyler *et al*., 2008). First, the assumption that we are working with a sample of unrelated individuals is particularly unrealistic in our scenario. Relatedness between individuals within and across the test and training sets impact GP accuracy (Habier *et al*., 2007; Pszczola *et al*., 2012; Dekkers *et al*., 2021), so the assumption of individuals being unrelated could bias the expected accuracy. Second, the assumption that no heterogeneous environmental effects are present in the system, implies that a phenotype *y*_*i*_ can simply be decomposed as *y*_*i*_ = *g*_*i*_ + *ε*_*i*_, where *ε*_*i*_ is an unstructured environmental residual, but in wild study systems the situation is usually more complicated. For example, one has little opportunity to control the environment, and thus has to account for environmental confounders as additional fixed or random effects in the model (Kruuk & Hadfield, 2007). In natural populations these environmental effects usually explain some of the phenotypic variance, and accounting for them is therefore important. In addition, the sampling design of a study might necessitate additional effects in the model, such as accounting for measurement error (Ponzi *et al*., 2018). In any case, it becomes unclear how to interpret the formula’s parameters in the presence of structured non-genetic effects in the model (see below), and certain steps in the derivations of formula (4) become invalid. In particular, it is assumed that the relation 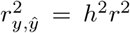 between *phenotypic* prediction accuracy *r*_*y,ŷ*_ and GP accuracy *r*. However, as shown in Appendix S3, 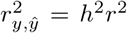 only holds when *y*_*i*_ = *g*_*i*_ + *ε*_*i*_, which is not true if we have other effects than *g*_*i*_ in the model.

Other limiting assumptions in the original derivation of (4) by Daetwyler et al. (2008) have been relaxed thanks to various generalizations (*e*.*g*., Goddard *et al*., 2011; Wray et al., 2013; Wientjes et al., 2015; Wray *et al*., 2019). For example, it is possible to account for the presence of relatedness by disentangling the genomic and pedigree-based sources of explanatory power (Dekkers *et al*., 2021).

However, the respective method requires fitting additional models that rely on information from pedigrees, whereas one of the main advantages of using genomic data in wild populations is that the laborious construction of pedigrees can be circumvented. Regarding the assumption of *y*_*i*_ = *g*_*i*_ + *ε*_*i*_, we note that it is common to pre-fit a mixed model with all non-genetic effects, and treating the identity effect from this non-genetic model as the new pseudo-phenotypes 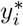 in a GP model (*e*.*g*., Dadousis *et al*., 2014; Hidalgo et al., 2016; Ashraf et al., 2022; Hunter et al., 2022; de Oliveira et al., 2023). By performing this pre-fitting step, the formula’s assumption of 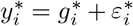 is true in the *second* model, given that we have accounted for all non-genetic effects. However, the variance explained in the first step is not accounted for in subsequent analyses, thus such a two-step approach could potentially bias the GP results (see *e*.*g*. Freckleton, 2002).

### Issues related to formula parameters

*Here we discuss issues related to the parameters of Equation (4), particularly M*_*e*_. A range of estimators exist for *M*_*e*_, each with somewhat differing assumptions (Brard & Ricard, 2015). The parameter *M*_*e*_ has, for instance, been estimated from effective population sizes, LD and genome length (Daetwyler *et al*., 2010; Goddard et al., 2011), from the variance of relatedness (Wientjes *et al*., 2015; Lee et al., 2017), or from rearranging Equation (4) and inserting for E (*r*) a realized 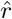 obtained in a previous model fitted to data from the same system (Dekkers *et al*., 2021). The problem is that it is not possible to know *a priori* which estimator for *M*_*e*_ is most appropriate in a particular scenario (Brard & Ricard, 2015, and see Appendix S3). Therefore, the only way to validate a given *M*_*e*_ estimate is to fit the model and compare the expected accuracy from (4) with the realized accuracy, but choosing an *M*_*e*_ estimator with this method would be circular reasoning. We also note that defining the heritability *h*^2^ in the presence of additional fixed and random effects is not always straightforward (see *e*.*g*. Wilson, 2008; de Villemereuil et al., 2018, and Appendix S3).

### Expected accuracy in across-population GP

In across-population GP, there are additional issues with current expected accuracy formulas, and the standard expected accuracy formula Equation (4) breaks down in the across-population case. Wientjes et al. (2015) therefore proposed scaling Equation (4) by the trait-specific genetic correlation between the two populations to correct for the decrease in accuracy when predicting across populations. However, estimating such genetic correlations is only possible if we have genotypic *and* phenotypic data from both populations, but if one has measured the phenotypes in the test population, then there is no reason to do a pure across-population prediction. In Appendix S3, we illustrate that even this scaling-approach failed at forecasting across-population GP accuracy for the house sparrow data. Furthermore, additional complications with expected accuracy formulas arise in across-population GP, such as the possibility that *h*^2^ is not equal in the two populations, or the presence of genotype-by-environment interactions, both of which can only be detected using phenotypes from both populations. Other commonly cited reasons for the decrease in accuracy such as population-differences in LD and MAFs (Wang *et al*., 2020), are also not accounted for by the formula in Equation (4).

## Population differentiation measures

The challenges with finding useful formulas to predict GP accuracy, especially across populations, reflect that the accuracy is impacted by underlying mechanisms that are not yet fully understood. To investigate which factors impacted across-population GP accuracies in our house sparrow applications, we estimated several measures of population differentiation between the training and test sets of each of the across-population models. As detailed below, we computed summaries of across-population genomic relatedness, a measure of population-differences in LD, a measure of population structure strength based on LD-inflation, and pairwise population genetic fixation indices (Wright’s *F*_*st*_, which reflect differences in allele frequencies). Similarly to the aforementioned expected accuracy formulas, these measures can be computed from genomic data only (as opposed to also requiring phenotypic measurements), which allows them to be used in an *a priori* consideration of whether GP is worthwhile. In Appendix S4, we additionally present estimates of trait-specific genetic correlations between training and test populations.

### Across-population genomic relatedness

Relatedness is expected to be a key factor in determining GP accuracy (Pszczola *et al*., 2012; Dekkers *et al*., 2021). For each model we therefore computed summary statistics on how related individuals in the training sets are to individuals in the test set. In other words, for a given model we considered a subset *G*_*ii*_*′*_,ac_ of entries of the GRM defined in Equation (2), such that individual *i* is in the training set and *i*′ is in the test set. To summarize these relatednesses, we computed the sample mean, which we denote 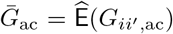. We also consider another summary statistic, the sample precision 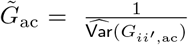, which has previously been considered important in determining GP accuracy (and indeed been employed as an estimator of the *M*_*e*_ parameter Wientjes *et al*., 2015; Lee et al., 2017).

### Population differences in linkage disequilibrium

The quality of GP models fundamentally relies on SNPs tagging quantitative trait loci (QTL) for the trait of interest (de los Campos *et al*., 2015). Thus, a critical type of difference between populations can lie in the patterns of LD within the respective training and test populations (de Roos *et al*., 2008). If there is a difference between the two populations in what loci are tagged by a given SNP, or the strength of that association, then the SNP-effects will in practice differ between the populations. To measure such discrepancies in LD between training and test sets, we first look at the difference in LD for a given pair of SNPs (*m, m*′)

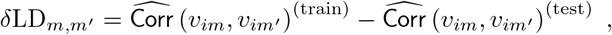

where *v*_*im*_ and *v*_*im*_*′* are the numbers of minor alleles that individual *i* has at *m* and *m*′, respectively. The superscripts indicate whether the Pearson correlations were computed using the individuals in the training or test set. The further the two populations have diverged, the more we expect the LD patterns in the populations to differ on average, so we consider the absolute mean difference in LD over all pairs

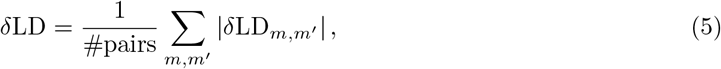

where we have taken the absolute value since it does not matter which direction *δ*LD_*m,m*_*′* changed. To reduce the computational burden, and since we are interested in how SNPs encode for nearby QTL, we limited our estimation of *δ*LD to SNP pairs located within 50 kilobases of each another (similar to Lundregan *et al*., 2018). Equation (5) captures how consistent the LD structure is between training and test populations, where large values of *δ*LD suggest that SNP pairs have different LD patterns in the two populations, which could undermine GP accuracy.

Because the presence of population structure is expected to inflate LD in a meta-population (Nei & Li, 1973), we also used LD-differences to measure the level of genetic differentiation between the training and test sets. Namely, we investigated how 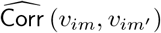 changes depending on whether population structure is taken into account or not. We define

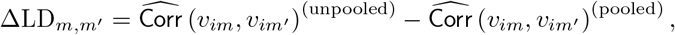

where the first term in ΔLD_*m,m*_*′* disregards the population structure by calculating 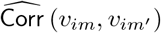 using all individuals in the combined training and test set as if they were drawn from a panmictic population, while the second term respects the distinction between training and test sets by accounting for population structure through pooling, that is, by properly aggregating the LD statistics from the training and test set. In other words, the numerator of 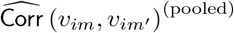 is the pooled covariance

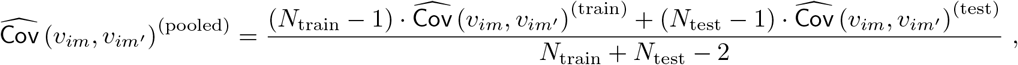

and the standard deviations in the denominator of 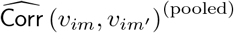 are treated similarly. As in Equation (5), we combine the ΔLD_*m,m*_*′* for different pairs of SNP using the absolute mean

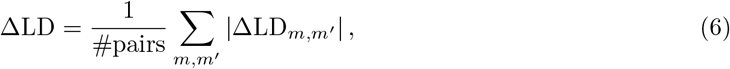

again restricting the calculation to SNP pairs within 50 kilobases. A high ΔLD indicates that LD was greatly inflated when not accounting for population structure, indicating large genetic differentiation between the training and test set, which in turn could impact GP accuracy negatively. To compute the correlations 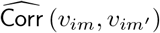 for the various sub-populations, we used `PLINK 1.9` (Chang *et al*., 2015).

### Pairwise population genetic fixation indices

*F*_*st*_ is a measure of genetic differentiation between populations, which essentially reflects differences in allele frequencies (Holsinger & Weir, 2009). Weir & Goudet (2017) present an expression for finding locus-specific estimates of *F*_*st*_ for pairs of populations, which we used to calculate the *F*_*st*_ between all pairs of training and test sets in our across-population GP models. We aggregated the locus-specific estimates using the method Weir & Goudet (2017) denote the “weighted” average, resulting in the *F*_*st*_ estimates

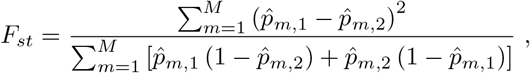

where 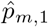 and 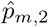 are the allele frequencies for SNP *m* in the training and test set, respectively. For each model, we estimated 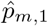 and 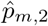 with `PLINK 1.9` (Chang *et al*., 2015).

## Results

### Within- and across-population GP accuracy

Using the observed accuracies from each fitted GP model, as defined in Equation (3), we found that within-population GP generally achieved higher accuracies than across-population GP (Figure 3). Despite the difference in training set sample sizes (see the Tables in Appendix S6), the within-population accuracies for a given trait were relatively similar in the three scenarios, with small differences in the mean accuracies relative to the variation. Conversely, the accuracies across populations in the Helgeland scenario were usually lower than in the within-populations scenarios, but highly variable, while the models from the southern and merged scenarios consistently performed more poorly in spite of the comparable training set sample sizes. The within-population accuracies were similar for the three traits, as were the across-population accuracies. However, the traits differed in the variability of the accuracies, both within and across populations, as accuracies for wing length were more variable than the accuracies for body mass, and the accuracies for tarsus length were the least variable (Figure 3).

**Figure 3:**
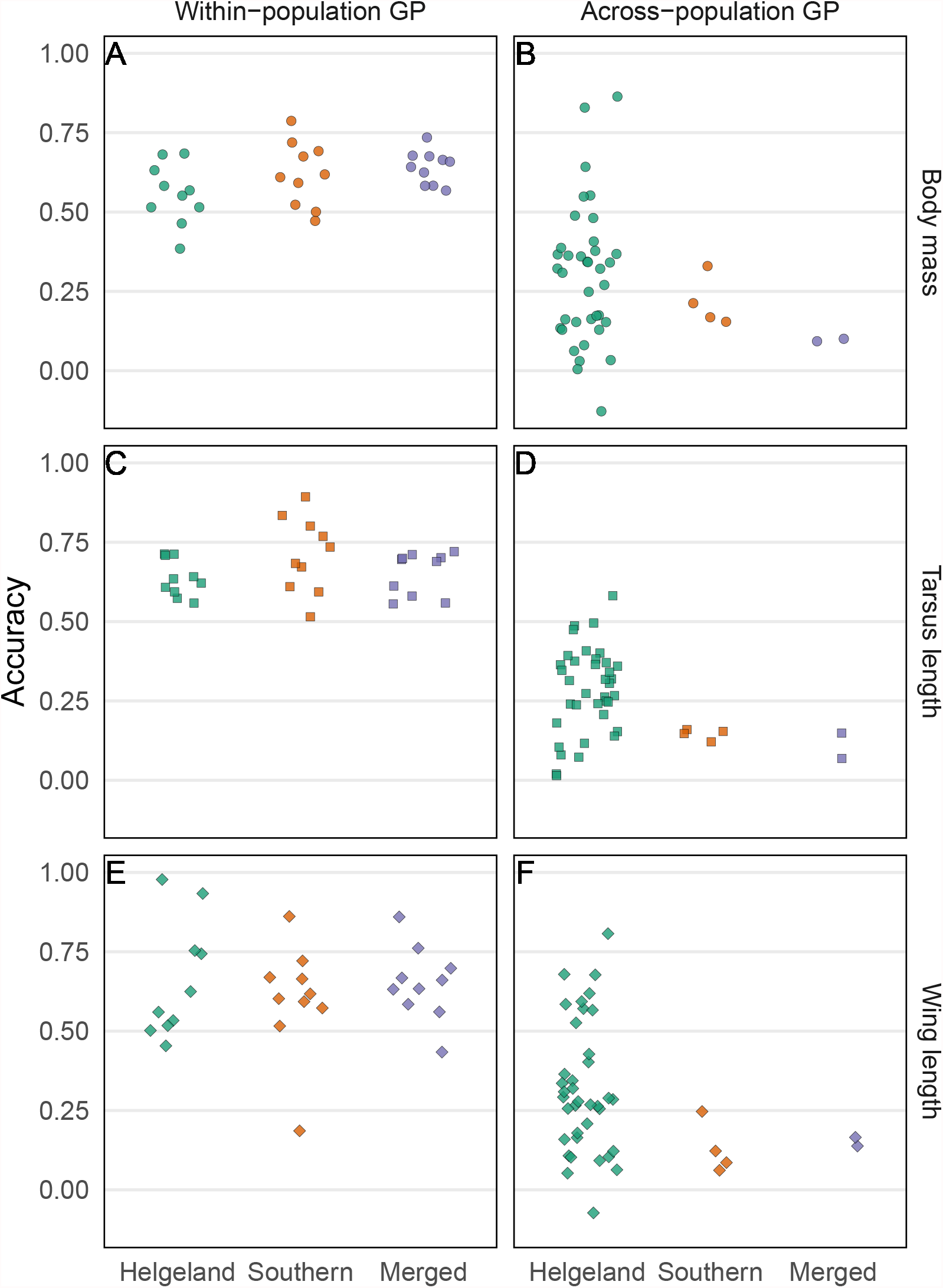
The accuracies from each GP model, as defined in Equation (3), for different scenarios and traits in the house sparrow systems. Models in panels **A** and **B** use body mass as the response, **C** and **D** use tarsus length and **E** and **F** use wing length. The left column of panels corresponds to within-population GP models, where each point is the accuracy for a single fold in the 10-fold CV. The right column of panels corresponds to the accuracies from the across-population GP models. Inside each panel, the points in green (left) correspond to accuracies from the Helgeland scenario, the points in orange (middle) correspond to accuracies from the southern scenario and the points in blue (right) correspond to accuracies from the merged scenario. The points are jittered horizontally to avoid overlap.

### Factors impacting across-population accuracy

We plotted the across-populations accuracies (*i*.*e*., the same ensemble of points as in the right column of panels in Figure 3) against various measures that potentially have an effect on accuracy (Figure 4). We found a slight trend for the across-population prediction accuracy to increase with training set size *N* (Figure 4A). However, accuracies were highly variable around this trend. For example, the models in Scenario 3 that were trained on the full Helgeland meta-population had the highest *N*, but also contained cases with some of the lowest accuracies. Higher mean relatedness 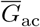 was associated with higher accuracy (Figure 4B). Notably, most of the across-population models had negative 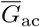, which can be interpreted as the average pair of across-population training and test individuals in those models sharing *fewer* alleles and thus being less related than would be expected at random. In other words, the individuals are – unsurprisingly – less related than if the training and test populations together formed a large panmictic population. The models formed two clusters of points, where the cluster with higher 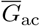 corresponded to every model that was tested on a non-farm island and trained on the remaining non-farm islands in the Helgeland system. This cluster contained across-population models that consistently yielded high prediction accuracy. Also the sample precision 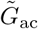 seemed associated with across-population accuracy, and the models with high accuracy had comparatively low 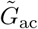 (Figure 4C). Note that the models from Scenario 2 (southern islands) were clustered together in Figure 4C, as were the models from Scenario 3 (merged), which had comparatively high 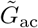 and low accuracy. Surprisingly, LD-differentiation measured by *δ*LD did not seem to impact across-population accuracy (Figure 4D), but we did find that across-population accuracies appeared to decrease with higher ΔLD (Figure 4E). Finally, across-population accuracy seemed not to be associated with pairwise *F*_*st*_ between the training and test populations (Figure 4F). The cluster of points with higher *F*_*st*_ correspond to all the models in the Helgeland scenario that had the test island Aldra, a highly inbred farm habitat island which was recently colonized by house sparrows (Billing *et al*., 2012). Note that there was a wide scatter of points around the smooth lines in all panels in Figure 4, indicating that none of the measures individually accounted for much variation in observed accuracy. In Appendix S4 we also investigated the impact of trait-specific genetic correlations between populations, and of heritability-related statistics, on across-population accuracy, while in Appendix S5 we performed multiple regression and variable importance analyses for the impact of the statistics in Figure 4, as well as E(*r*), on across-population accuracy. In brief, Appendix S5 shows that *F*_*st*_ and *δ*LD are highly correlated, as are ΔLD and E(*r*), and that the parameters in Figure 4 explained less than half of the variance in across-population accuracy, but more variance than was explained by E(*r*). Appendix S6 contains detailed tables with results from all fitted GP models.

**Figure 4:**
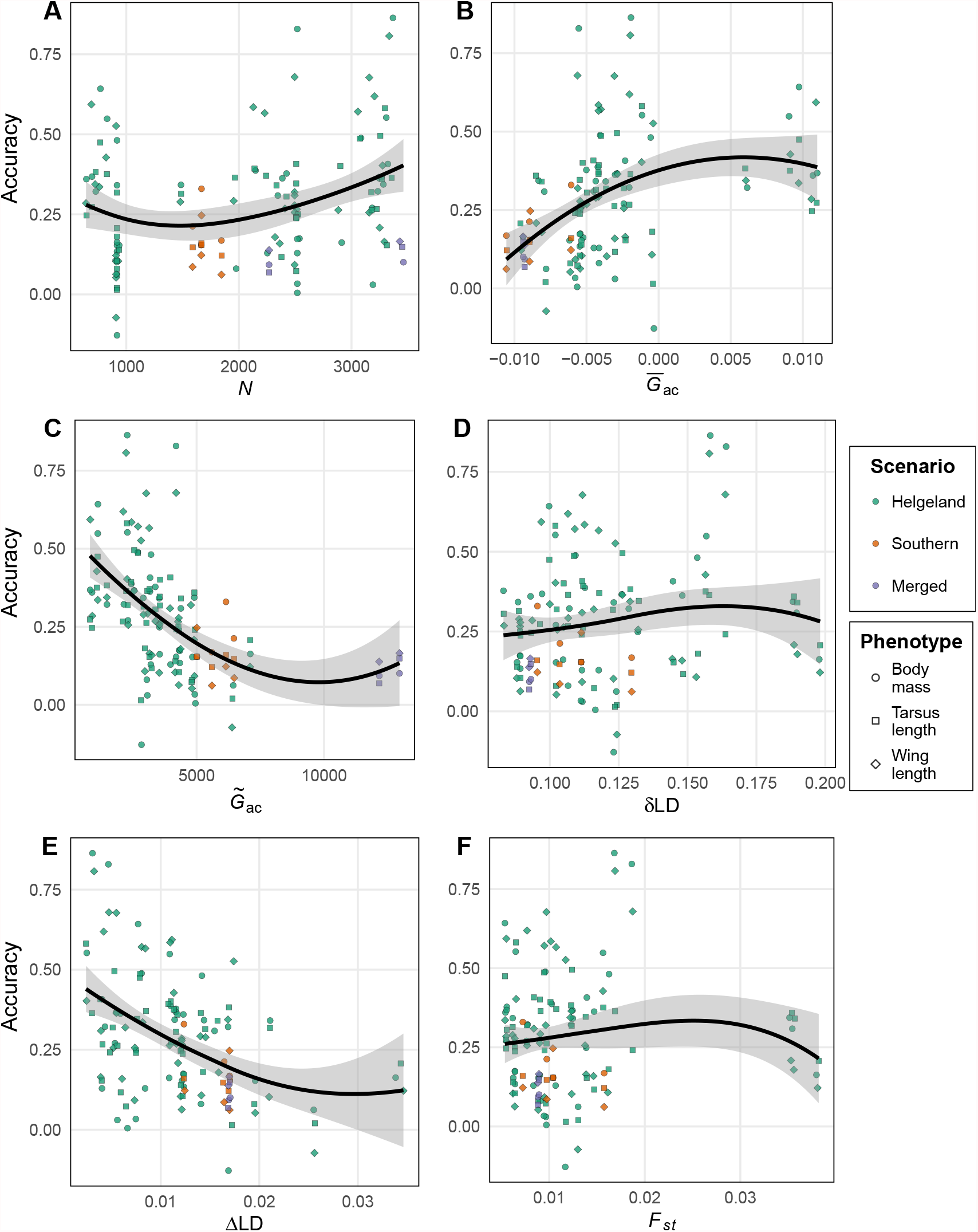
Scatter plots of across-population prediction accuracies for all scenarios and traits, plotted against various statistics. The statistics on the *x*-axis correspond to the number of individuals *N* in the training set (**A**), the mean relatedness 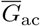 between individuals in the training and test set (**B**), the precision 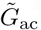 of relatedness between individuals in the training and test set (**C**), *δ*LD, a measure of LD-differentiation between the populations (**D**), ΔLD, a measure of population structure between the training and test set (**E**), and the population-pairwise fixation index *F*_*st*_ (**F**). The smoothing lines are produced with local regression.

## Discussion

In this article we applied GP models to morphological phenotypes in wild populations of house sparrows. As the use of GP in wild populations is still in its infancy, our results have important implications for future applications. We demonstrated that GP achieves relatively high accuracies when applied within populations. GP is thus a promising tool for the field of evolutionary ecology, as it lets us effectively predict breeding values for non-phenotyped individuals. However, our results illustrate that across-population applications should currently be carried out with caution, as existing GP approaches are considerably less accurate when applied across populations.

Since our understanding of the exact mechanisms that determine across-population GP accuracy is limited, we empirically investigated the impact of various population differentiation statistics. Several measures appear to be important to across-population accuracy, including the mean 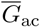 and precision 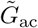 of the relatedness between individuals in the two populations and the population structure measure ΔLD. Based on our results, we cannot recommend applying GP across populations unless 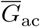 is high, 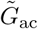 is low, and/or ΔLD is low, as these situations are most consistently associated with high accuracies. However, if there only is one training and one test population, it might be unclear what acceptable values for the investigated statistics are. And even for given values of the population differentiation measures, across-population accuracy is usually highly unpredictable (*i*.*e*., the *y*-axes of Figure 4 are highly variable for most values along the *x*-axes). A major problem is that we still lack an *a priori* understanding of when across-population GP performs well or not, as we do not have a reliable formula to predict GP accuracy in a given scenario. A fuller and more mechanistic understanding of these underlying processes is required to reliably apply GP across-populations.

Our finding that existing GP models perform better within populations raises the question of what to do when only limited phenotypic data are available from a population of interest, but we have access to more measurements from other, distant populations. Restricting ourselves to only using GP within population would make populations that are small and isolated – the populations with the highest risk of extinction – especially challenging applications, as sample sizes will necessarily be small. This potentially problematic issue may be counteracted by the other consequences of populations being smaller and more isolated: they will have higher levels of relatedness and LD, meaning that GP is expected to be more accurate. In any case, if one has a few phenotypes from the focal population, but also a large number of measurements from *other* populations, then the training set could in practice be “augmented” with data from these other populations. This approach could be beneficial in terms of increasing the sample size of the training set, but also risks degrading model performance due to the various factors decreasing across-population accuracy. We did not consider such training set augmentation here, and instead focused on “pure” across-population GP, where the the training and test data are from entirely disjoint populations. Utilizing data from both the population of interest and other populations is a more general problem which comes down to maximizing GP accuracy by using the available data in the best possible way. This gives rise to the idea of “training set optimization” methods, where one aims to determine which individuals should be included in the training set to optimize the accuracy in a given test set (*e*.*g*., Rio *et al*., 2021). Investigating such optimization approaches in wild populations would be an interesting focus for a future study, in particular in the context of conservation efforts.

One potential weakness of our analysis is that we only considered relatedness-based GP models (*i*.*e*., genomic animal models, G-BLUP). These models were a natural choice since they are considered the baseline GP model against which other models are often compared (*e*.*g*. Ashraf *et al*., 2022; Aspheim *et al*., 2024). Other GP methods could potentially improve general accuracy (*e*.*g*., BayesR, see Yin *et al*., 2022), or better deal with the problematic aspects of across-population GP. However, accuracy gains within populations for other GP compared to G-BLUP are usually marginal for highly polygenic traits like the morphological phenotypes we investigated (see Ashraf *et al*., 2022). It is worth pointing out that we deliberately made no special effort to account for the models being applied across populations, which explicitly violates the assumptions of G-BLUP regarding lack of population structure (see *e*.*g*. Wolak & Reid, 2017). Instead, we straightforwardly applied the models as one would in the within-population case, rather than accounting for, for example, population-differences in allele frequencies (see *e*.*g*. Wientjes *et al*., 2017). Thus, our study also investigated potential negative consequences of breaking these assumptions. Exploring alternative GP models, such as marker-regression approaches (*e*.*g*., Yin *et al*., 2022; Aspheim et al., 2024), or methods which explicitly incorporate population structure (*e*.*g*., Rio *et al*., 2020; Aase *et al*., 2022), could improve across-population GP, and should be examined in future studies.

By considering empirical data from a wild population, we ensured that the range of situations that we investigated were both realistic and ecologically relevant. Notably, the study covers the full spectrum of possible outcomes from across-population GP, as we notably found observed accuracies throughout the interval [0, 1]. On the other hand, our approach also limited which dimensions of the problem we were able to investigate. For instance, the traits we looked at are all continuous, highly polygenic and have similar heritabilities (Silva *et al*., 2017, and see Appendix S6). Future research on across-population GP in wild populations could therefore benefit from considering a broader range of traits and incorporating simulation studies with known ground truths.

In conclusion, GP is an emerging tool in evolutionary ecology, with the potential to effectively predict breeding values of genotyped, but not necessarily phenotyped, individuals. While GP is reliable within single (meta-)populations, more work is needed to better understand and utilize the promise of GP across populations. In particular, we need to understand the factors that enhance or limit prediction accuracy, and whether it is possible to bypass these limitations. This would allow us to fully harness the tremendous potential of genomic data using GP, in order to understand evolutionary processes across space and time and, ultimately, help address the biodiversity crisis.

## Acknowledgments

We thank the many researchers, students, and fieldworkers who helped in collecting the empirical data on house sparrows, and laboratory technicians for assistance with laboratory analyses. This work was possible thanks to generous funding from the Department of Mathematical Sciences at Norwegian University of Science and Technology, as well as by grants from the European Research Council (grant number 101169862), and the Research Council of Norway (project numbers 274930 and 302619). This work was also partly supported by the Research Council of Norway through its Centres of Excellence funding scheme (project number 223257). Genotyping on the custom house sparrow Axiom 200K and 70K SNP arrays was carried out at CIGENE, Norwegian University of Life Sciences, Norway. The computations were performed on resources provided by the NTNU IDUN/EPIC computing cluster (Själander *et al*., 2019). We thank Jarrod D. Hadfield for the useful discussion which inspired the LD-measures in the analysis.

## Conflict of Interest statement

The authors have no conflicts of interest to declare.

### Authors’ contributions

KA, SM and HJ conceived the idea for the study. KA performed and implemented all analyses, with support from SM, HJ and HB. HJ and HB provided the sparrow data. KA wrote the manuscript, with support from SM and HJ. KA, SM and HJ edited the manuscript. All authors read and approved the final manuscript.

## Data availability

The data underlying this article are subject to an embargo of 12 months from the publication date of the article. Once the embargo expires the data will be available upon reasonable request. The code used to generate the results is available here.

## Ethical Statement

The research done on house sparrows was carried out in accordance with permits from the Norwegian Food Safety Authority and the Norwegian Bird Ringing Centre at Stavanger Museum, Norway.

The first author would like to note that citation of literature from the field of animal breeding is not an endorsement of the practices of the meat, dairy or egg industries. Conversely, the first author affirms the intrinsic moral value of non-human animals, and thus opposes the unnecessary and unimaginable suffering experienced by the victims of industrial animal agriculture.

