## Supporting Information for "How accurate is genomic prediction across wild populations?"

### S1: Genomic animal models

To estimate variance components and heritabilities in subpopulations, we fitted 9 genomic animal models, one for each trait in each scenario. These models were of the same form as the GP models described in the main text, except all available data for a given trait/scenario was used to train the models (*i.e.*, there was no test set). The same fixed and random effects as in Equation (1) in the main text were included, the GRM was constructed with the same procedure, and the same priors were used in `R-INLA`.

By sampling from the posteriors of these respective animal models, we estimated heritabilities and variance components in subsets of the data. Additive genetic variances and heritabilities were estimated as the posterior means

$$\hat{\sigma}^2_{\mathrm{G}} = \frac{1}{S}\sum_{s=1}^{S}\hat{\sigma}^2_{\mathrm{G},(s)} = \frac{1}{S}\sum_{s=1}^{S}\widehat{\mathsf{Var}}\left(g_i^{(s)}\right) \ , \quad \hat{h}^2_{\mathrm{AM}} = \frac{1}{S}\sum_{s=1}^{S} h^2_{(s)} = \frac{1}{S}\sum_{s=1}^{S}\frac{\widehat{\mathsf{Var}}\left(g_i^{(s)}\right)}{\widehat{\mathsf{Var}}\left(\tilde{y}_{ij}^{(s)}\right)} \ , \tag{S1}$$

respectively, where $\hat{\sigma}^2_{\mathrm{G}}$ is the estimated additive genetic variance, $S = 10\ 000$ is the chosen number of posterior samples, the index $(s)$ indicates the $s^{\mathrm{th}}$ posterior sample, $\hat{\sigma}^2_{\mathrm{G},(s)}$ is a posterior sample of the additive genetic variance, $g_i^{(s)}$ is a posterior sample of the genetic value, $\hat{h}^2_{\mathrm{AM}}$ is the estimated heritability, $h^2_{(s)}$ is a posterior sample of the heritability, $\tilde{y}_{ij}^{(s)}$ is a posterior sample of the linear predictor from the model excluding $\varepsilon_{ijk}$ (see Equation (1) in the main text) and $\widehat{\mathsf{Var}}$ denotes sample variance. We use $\tilde{y}_{ij}$ (rather than the data $y_{ijk}$) to exclude the variance caused by the measurement error $\varepsilon_{ijk}$ from the denominator, since it is not a biologically relevant source of variation (see de Villemereuil *et al.*, 2018). This is in contrast to the accuracy scaling factor $\lambda^2$ used in the main text, which *does* use the sample variance in the data $y_{ijk}$ in its denominator.

Here, $\widehat{\mathsf{Var}}$ denotes sample variance taken over the individuals $i$ in a given subset of the individuals in a given scenario (Sorensen *et al.*, 2001; Lara *et al.*, 2022). This lets us find additive genetic variances and heritabilities in subsets of the data, for example in the test set of a given GP model. We use the estimates $\hat{\sigma}^2_{\mathrm{G}}$ of additive genetic variance in a given test set to find the proper scaling factor $\lambda$ for GP accuracy (see main text and Appendix S2), and the estimates $\hat{h}^2_{\mathrm{AM}}$ are used in Appendix S3 in the context of formulas for expected accuracy, and are also used in Appendix S4 in comparisons the test set heritabilities derived from the GP models in the main text. Results for the posterior statistics for the genomic animal are shown in Appendix S6.

### S2: GP accuracy metrics

#### Derivation of accuracy scaling factor

Here we show that our accuracy measure (Equation (3) in the main text) is equivalent to $r = \mathsf{Corr}\left(g_i, \hat{g}_i\right)$. We assume that among all the model terms in $y_{ijk}$, the only one that covaries with $\hat{g}_i$ is $g_i$, which essentially means that our model is correctly specified, and so we have accounted for all confounding factors. Thus, we can write

$$
\begin{aligned}
\mathsf{Corr}\left(y_{ijk}, \hat{g}_i\right) &= \frac{\mathsf{Cov}\left(y_{ijk},\ \hat{g}_i\right)}{\sqrt{\mathsf{Var}\left(y_{ijk}\right)\ \mathsf{Var}\left(\hat{g}_i\right)}} \\
&= \frac{\mathsf{Cov}\left(\ldots + g_i + \ldots,\ \hat{g}_i\right)}{\sqrt{\mathsf{Var}\left(y_{ijk}\right)\ \mathsf{Var}\left(\hat{g}_i\right)}} \\
&= \frac{\mathsf{Cov}\left(g_i,\ \hat{g}_i\right)}{\sqrt{\mathsf{Var}\left(y_{ijk}\right)\ \mathsf{Var}\left(\hat{g}_i\right)}} \\
&= \frac{\mathsf{Cov}\left(g_i,\ \hat{g}_i\right)}{\sqrt{\mathsf{Var}\left(y_{ijk}\right)\ \mathsf{Var}\left(\hat{g}_i\right)}} \cdot \sqrt{\frac{\mathsf{Var}(g_i)}{\mathsf{Var}(g_i)}} \\
&= \frac{\mathsf{Cov}\left(g_i,\ \hat{g}_i\right)}{\sqrt{\mathsf{Var}\left(g_i\right)\ \mathsf{Var}\left(\hat{g}_i\right)}} \cdot \sqrt{\frac{\mathsf{Var}(g_i)}{\mathsf{Var}(y_{ijk})}} \\
&= \mathsf{Corr}\left(g_i,\ \hat{g}_i\right) \cdot \lambda\ , \\
&= r \cdot \lambda\ ,
\end{aligned}
$$

where $\lambda = \sqrt{\frac{\mathsf{Var}(g_i)}{\mathsf{Var}(y_{ijk})}}$ as defined in the main text. In other words, we define $\lambda^2$ as the variance ratio between the genetic values and the phenotypic measurements, *including the measurement error*. As discussed by Wilson (2008) and de Villemereuil *et al.* (2018), the proper definition of heritability should not involve non-biological sources of variation such as measurement error in the denominator. Thus, we should not write $\lambda = h$ as is commonly done in the literature, as $\lambda$ will only be equal to the square root of the heritability under a naïve definition of heritability.

#### Uncertainty in GP accuracy

Assessing the uncertainty in the GP accuracy $\hat{r}$ is straightforward in our within-population models. We can use $\hat{\sigma}_{\mathrm{CV}}$ to estimate the uncertainty, where $\hat{\sigma}_{\mathrm{CV}}$ is the sample standard deviation over the 10 accuracies obtained from a given 10-fold cross-validation. However, for the across-population GP models, we only get a single point estimate of accuracy per model. One might think that it would have been enough to take joint posterior samples of the genetic value vector $\boldsymbol{g}$ from a given across-population GP model, and use these each of these samples in Equation (3) in the main text, but doing this will severely underestimate the accuracy compared to using $\hat{g}_i$. The joint posterior of the GP models give us access to the uncertainty in $g_i$, but what we are interested in is the uncertainty in the point estimate $\hat{g}_i$. Thus, we need multiple samples of $\hat{g}_i$, which implies a need for refitting the model.

Here we suggest a procedure that involves resampling from the joint posterior of the fitted models, which is similar to a parametric bootstrap. For each GP model, we resampled 100 new response vectors $\boldsymbol{y}^{(b)}(b = 1, \ldots, 100)$ from the fitted joint posterior of the model. For each new resampled response vector $\boldsymbol{y}^{(b)}$ we then refitted the model and retained the accuracy of the genetic values obtained from the refitted model, $\hat{r}^{(b)}$, calculated using Equation (3) with $g_i$ and $y_{ijk}$ replaced by $g_i^{(b)}$ and $y_{ijk}^{(b)}$, respectively. This procedure thus produces 100 replicates $\hat{r}^{(b)}$ of the accuracy $\hat{r}$ of a given GP model, and we can take the sample standard deviation of these replicates to estimate the uncertainty of $\hat{r}$, or look at other summary statistics such as quantiles.

The resampling and refitting procedure has the obvious downsize of being computationally cumbersome, as it requires fitting each GP a large number of additional times, *e.g.*, 100 times. Because of the large computational requirements, we only estimated the uncertainty in a subset of the across-population GP models, namely the models for tarsus length in Scenario 1 (Figure S1), as well as the all models in Scenario 2 (Figure S2).

In Figure S3 we demonstrate that the uncertainties from the resampling method are well-calibrated. That is, for a given probability level $\pi \in (0, 1)$, we compute the empirical coverage as the proportion of models for which the realized accuracy $\hat{r}$ falls within the central $\pi$-quantile range $\left[0.5 - \frac{\pi}{2}, 0.5 + \frac{\pi}{2}\right]$ of the corresponding resampled accuracies $\hat{r}^{(b)}$. We find a close correspondence between this empirical proportion and its theoretical expectation (which is equal to $\pi$), indicating a well-calibrated uncertainty measure.

### S3: Analysis of formulas for expected accuracy

Here we investigate the efficacy of the expected accuracy formula

$$\mathsf{E}(r) \approx \sqrt{\frac{1}{1 + \frac{M_e}{Nh^2}}}\ , \tag{S2}$$

given as equation (4) in the main text, and the more general version

$$\mathsf{E}(r)^2 = \frac{\frac{h_M^2}{h^2}}{1 + \frac{M_e}{N \cdot h_M^2} \cdot \left(1 - h^2 \mathsf{E}(r)^2\right)}\ , \tag{S3}$$

given by Wray *et al.* (2013). Here $\mathsf{E}(r)$, $h^2$, $N$ and $M_e$ are as given in the main text, while $h_M^2$ is the heritability captured by $M$ genetic markers. By differentiating between $h^2$ and $h_M^2$, the generalized formula (S3) allows for the possibility that one has not included enough markers to capture all the

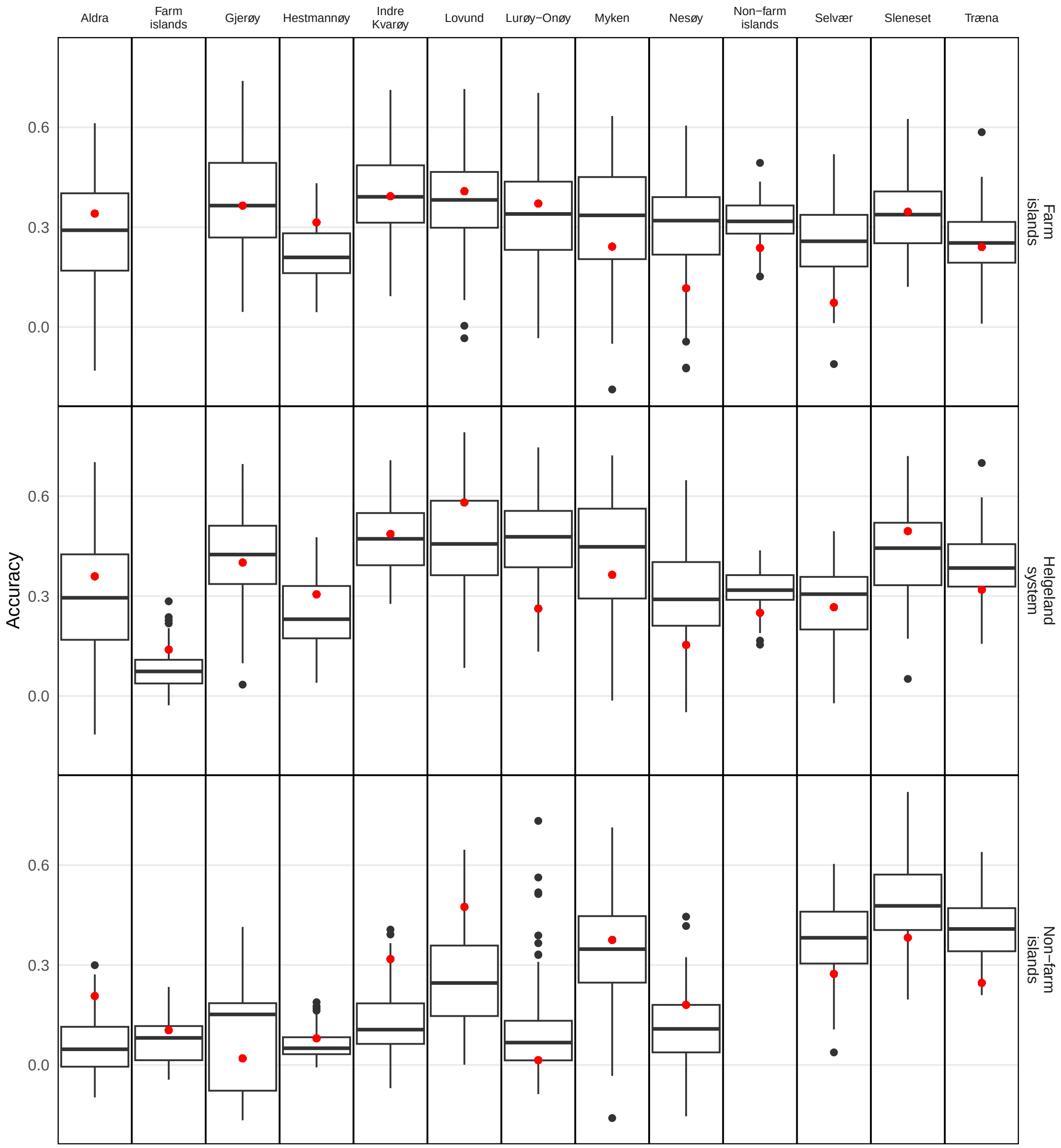


Figure S1: Accuracy in the across-population GP models for tarsus length. Columns correspond to test sets, and rows correspond to training sets, excluding the test island(s) if relevant. The red dots are the point estimates of accuracy obtained from the original GP model and presented in the main text, while the box plots are made from the accuracies from the refitted models.

additive genetic effects, that is, to tag all the QTL of the polygenic trait. And, in contrast to formula (S2), the generalization (S3) contains the factor $\left(1 - h^2\mathsf{E}(r)^2\right)$ in the denominator, which makes the equation quadratic in $\mathsf{E}(r)^2$. By assuming $h^2 = h^2_M$ (*i.e.*, *marker saturation*, the markers capture the full heritability) and $h^2\mathsf{E}(r)^2 \approx 0$ we retrieve the original formula (S2).

As we discussed in the main text, several questions arise about which choices should be made when applying the expected accuracy formulas such as (S2) and (S3). Here we compare the performance of

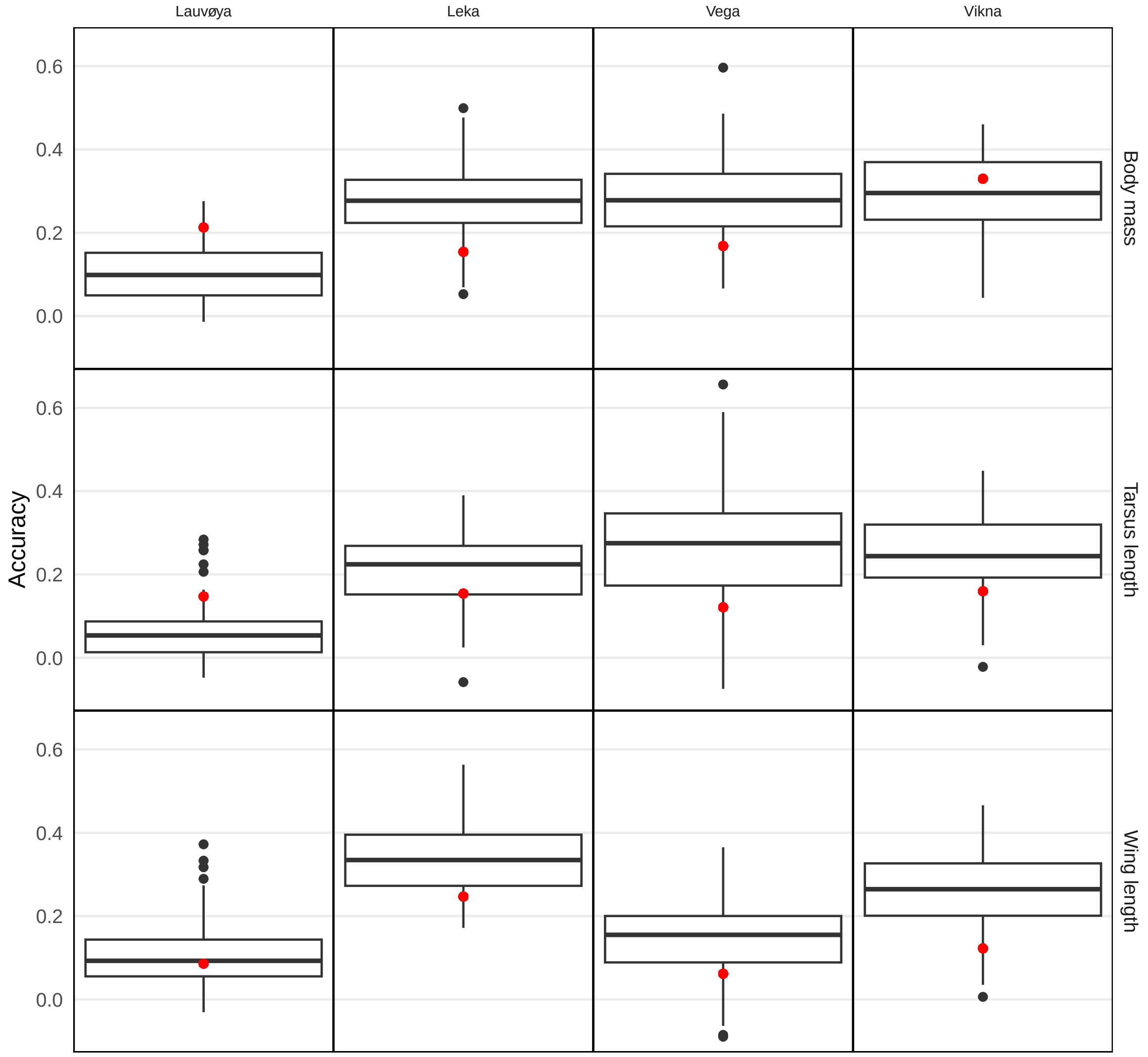


Figure S2: Accuracy in the across-population GP models for Scenario 2. Columns correspond to test sets, and rows correspond to phenotypes. The training sets are the sparrows from the southern islands, excluding the test island. The red dots are the point estimates of accuracy obtained from the original GP model and presented in the main text, while the box plots are made from the accuracies from the refitted models.

the expected accuracy formulas under combinations of choices related to $M_e$, heritability $h^2$ and which formula to use. The set of choices we considered are detailed below.

##### Choice of $M_e$ estimator

We investigated 7 different estimators of $M_e$: the five estimators denoted $M_{e1}$ through $M_{e5}$ by Brard & Ricard (2015):

$$M_{e1} = \frac{2N_eL}{\log(4N_eL)},$$

$$M_{e2} = \frac{2N_eL}{\log\left(2N_e\frac{L}{N_{\text{chr}}}\right)},$$

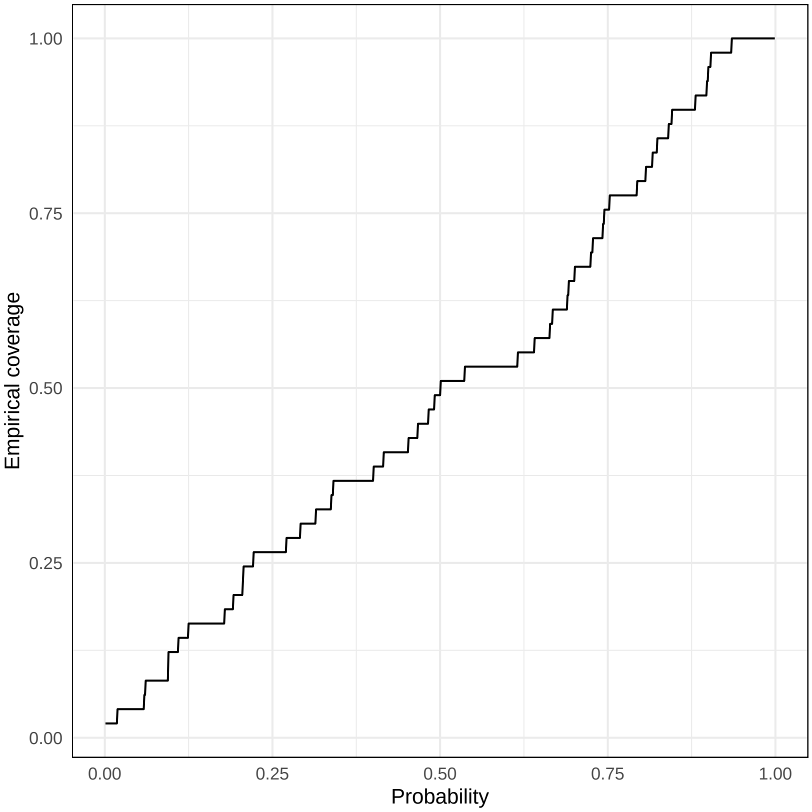


Figure S3: Empirical coverage of the resampling method. For each probability level ($x$-axis) we compute the proportion of models ($y$-axis) whose true accuracies $\hat{r}$ fall within the central $\pi$-quantile range of their corresponding resampled accuracies $\hat{r}^{(b)}$.

$$M_{e3} = \frac{2N_e L}{\log\left(N_e \frac{L}{N_c}\right)},$$

$$M_{e4} = 2N_e L,$$

$$M_{e5} = 4N_e L,$$

as well as two estimators which Lee *et al.* (2017) denote as equation (2) and (5), that we respectively label $M_{e6}$ and $M_{e7}$:

$$M_{e6} = \frac{N_{\text{chr}}}{\frac{\ln\left(2N_e \frac{L}{N_{\text{chr}}}+1\right)+2N_e \frac{L}{N_{\text{chr}}}\left(\ln\left(2N_e \frac{L}{N_{\text{chr}}}+1\right)-1\right)}{4N_e^2 \frac{L^2}{N_{\text{chr}}^2}} + \frac{1}{3N_e}\left(N_{\text{chr}}-1\right)},$$

$$M_{e7} = \tilde{G}_{\text{ac}} = \frac{1}{\widehat{\mathsf{Var}}(G_{ii',\text{ac}})},$$

where $\tilde{G}_{\text{ac}}$ is as defined in the main text. Estimators $M_{e1}$ through $M_{e6}$ are based on the effective population size $N_e$, genome length $L$ in Morgans and the number of chromosomes $N_{\text{chr}}$. These estimators use $N_e$, a population parameter, which thus implicitly also defines $M_e$ as a population parameter.

Conversely, the last estimator $M_{e7}$ can be considered as taking a model-specific view of $M_e$, as it considers the genomic relationships between the individuals in the training and test sets for the model.

For each model in our study we found 7 estimates of $M_e$ by using the estimators $M_{e1}$ through $M_{e7}$. To estimate $N_e$ we used the software `currentNe` (Santiago *et al.*, 2024), whereas the parameters related to the house sparrow genome parameters were taken from Hagen *et al.* (2020). We always estimated $N_e$ for the individuals in the *combined* training and test sets, including when considering the across-population models.

#### Heritability-related choices

We also investigated assumptions related to how heritability should be defined when applying expected accuracy formulas. We looked at the following heritability-related choices when using $h^2$ in the expected accuracy formulas (S2) and (S3):

1. Should we assume marker saturation, *i.e.*, should we assume that $h^2_M = h^2$ or not? When assuming marker assumption we set $h^2_M = h^2 = \hat{h}^2$. When *not* assuming marker saturation we set $h^2_M = \hat{h}^2$ and estimated $h^2$ in each model using the formula $\frac{h^2_M}{h^2} = \frac{M}{M+M_e}$ (Goddard *et al.*, 2011), where $M$ is the number of SNP markers and $M_e$ is one of the 7 estimates outlined in the previous subsection.

2. Which subset of data should be used to estimate $h^2$? That is, should we use the heritability among the individuals in the combined training and test set, or just the individual in the test set? These heritabilities are assumed to be equal when expected accuracy formulas are derived (*e.g.*, Wientjes *et al.*, 2015). However, this question is relevant in the context of across-population GP, where the heritability of the training and test sets could be notably different. It would be problematic if the formula only performs well when using the heritability in the test set, since we would need the phenotypes in the test set to estimate this heritability, which, as explained in the main text, defeats the purpose of a formula for expected accuracy. Thus, when applying equation (S1) from Appendix S1 to estimate heritabilities, we computed $\hat{h}^2$ using samples $g_i^{(s)}$ and $\tilde{y}_{ijk}^{(s)}$ either from all individuals or just the subset of individuals in the test set.

#### Quadratic formula

Finally, we investigated whether one should include the term $\left(1 - h^2 \mathsf{E}\left(r\right)^2\right)$ in the denominator of the formula, as in (S3), or not, as in (S2). That is, should one solve a quadratic equation for $\mathsf{E}\left(r\right)^2$, or assume that $h^2\mathsf{E}(r)^2 \approx 0$?

#### Gentic correlations

(Wientjes *et al.*, 2015) suggest that in across-population GP, formulas for expected accuracy should be scaled by estimated genetic correlations between the training and test populations. We investigated whether this scaling improved the performance of the expected accuracy formulas for our across-population models, by using estimates of genetic correlation from Appendix S4.

#### Analysis of formula-choices

We now investigate how well the expected accuracy formulas perform depending on the choices outlined above. As we considered 7 options for $M_e$, and three additional binary choices (marker saturation, choice of subset for $h^2$, quadratic formula or not), we investigated $7 \cdot 2 \cdot 2 \cdot 2 = 56$ combinations in the within-population case. In the across-population case we also checked whether scaling by genetic correlations improved fits, adding another binary choice, and thus increasing the number of combinations to $56 \cdot 2 = 112$. For each combination $c \in \{1, \dots, 56\}$ in the within-population case and $c \in \{1, \dots, 112\}$ in the across-population case, we evaluated how well the respective expected accuracy formula performed by computing the RMSE $\delta_c^{\mathcal{M}}$ between realized and expected accuracies;

$$\delta_c^{\mathcal{M}} = \sqrt{\frac{1}{|\mathcal{M}|} \sum_{\mathfrak{m} \in \mathcal{M}} (r_{\mathfrak{m}} - \mathsf{E}(r)_{\mathfrak{m},c})^2} \ ,$$

where $\mathcal{M}$ is a set of models, $r_{\mathfrak{m}}$ is the realized accuracy in model $\mathfrak{m}$ and $\mathsf{E}(r)_{\mathfrak{m},c}$ is the expected accuracy for model $\mathfrak{m}$ when using the choices in combination $c$. We computed $\delta_c^{\mathcal{M}_{\text{within}}}$ where $\mathcal{M}_{\text{within}}$ is the set of within-population models (with $|\mathcal{M}_{\text{within}}| = 90$), and $\delta_c^{\mathcal{M}_{\text{across}}}$ where $\mathcal{M}_{\text{across}}$ is the set of across-population models (with $|\mathcal{M}_{\text{across}}| = 129$).

Table S1: Effect sizes, standard errors and associated $p$-values from the linear models investigating how different choices relating to expected accuracy formulas impact the expected accuracies' RMSE with regards to realized accuracy in the witihn-population case.

| | Estimate | Std. Error | p-value |
|---|---|---|---|
| Intercept | 0.1294 | 0.0014 | 0.0000 |
| $X_{M_{e1}}$ | 0.0180 | 0.0016 | 0.0000 |
| $X_{M_{e2}}$ | 0.0129 | 0.0016 | 0.0000 |
| $X_{M_{e3}}$ | 0.0192 | 0.0016 | 0.0000 |
| $X_{M_{e4}}$ | 0.3217 | 0.0016 | 0.0000 |
| $X_{M_{e5}}$ | 0.3784 | 0.0016 | 0.0000 |
| $X_{M_{e6}}$ | 0.0014 | 0.0016 | 0.3789 |
| $X_{h^2=h^2_M}$ | 0.0014 | 0.0009 | 0.1242 |
| $X_{h^2_{\text{test}}}$ | 0.0010 | 0.0009 | 0.2381 |
| $X_{r^2h^2\neq 0}$ | -0.0035 | 0.0009 | 0.0002 |

We fitted two linear regression models with $\delta_c^{\mathcal{M}_{\text{within}}}$ and $\delta_c^{\mathcal{M}_{\text{across}}}$ as the responses, respectively, and dummy variables $X_\bullet \in \{0, 1\}$ encoding for the different combinations as covariates. $X_{M_{e1}}$ through

Table S2: Effect sizes, standard errors and associated $p$-values from the linear models investigating how different choices relating to expected accuracy formulas impact the expected accuracies' RMSE with regards to realized accuracy in the across-population case.

| | Estimate | Std. Error | p-value |
|---|---|---|---|
| Intercept | 0.2015 | 0.0107 | 0.0000 |
| $X_{M_{e1}}$ | 0.0423 | 0.0121 | 0.0007 |
| $X_{M_{e2}}$ | 0.0450 | 0.0121 | 0.0003 |
| $X_{M_{e3}}$ | 0.0418 | 0.0121 | 0.0008 |
| $X_{M_{e4}}$ | 0.0539 | 0.0121 | 0.0000 |
| $X_{M_{e5}}$ | 0.0767 | 0.0121 | 0.0000 |
| $X_{M_{e6}}$ | 0.0535 | 0.0121 | 0.0000 |
| $X_{h^2=h^2_M}$ | -0.0006 | 0.0065 | 0.9230 |
| $X_{h^2_{\text{test}}}$ | -0.0007 | 0.0065 | 0.9167 |
| $X_{r^2h^2\neq 0}$ | 0.0023 | 0.0065 | 0.7283 |
| $X_{\text{gc}}$ | -0.0168 | 0.0065 | 0.0106 |

$X_{M_{e6}}$ encode for the respective choice of $M_e$ estimator (with $M_{e7}$ being the reference category), $X_{h^2=h^2_M} = 1$ when marker saturation ($h^2 = h^2_M$) was assumed, $X_{h^2_{\text{test}}} = 1$ when the heritability in the test was used and $X_{r^2h^2\neq 0} = 1$ for combinations where we assume $r^2h^2 \neq 0$. $X_{\text{gc}} = 1$ when the formula for expected accuracy was scaled by estimated genetic correlations. We can thus investigate which choices (conditional on the other choices) are associated to a lower RMSE $\delta_c^{\mathcal{M}}$, *i.e.*, a better fit between the realized and expected accuracies. The results from the linear models, as shown in Tables S1 and S2, are that:

- We found strong evidence that $M_{e7}$ performed better (*i.e.*, was associated with lower RMSE) than $M_{e1}$ through $M_{e5}$, for both within-population and across-population models. For within-population models there was no evidence that $M_{e6}$ and $M_{e7}$ performed differently, but there was strong evidence that $M_{e7}$ performed better than $M_{e6}$ for across-population models. Lee *et al.* (2017) states that $M_{e6}$ and $M_{e7}$ are equivalent, but interestingly this appears to only holds in the within-population case - in the across-population case $M_{e6}$ is among the worst but $M_{e7}$ is the best.

- $M_{e4}$ and $M_{e5}$ are drastically worse than $M_{e7}$ in the within-case, with the overall largest predicted increase in RMSE. However, the increase in RMSE is much smaller for these estimators in the across-case, indicating that what might be a reasonable choice in one scenario could be disastrous in another.

- There was weak evidence that assuming marker saturation ($h^2 = h^2_M$) increased the RMSE for within-population models, but no evidence for across-population models.

- There is no evidence that using the heritability in the test set degraded performance of the expected accuracy formula compared to using the heritability in the combined training and test sets.

- We found strong evidence that the quadratic version of the expected accuracy formula (*i.e.*, including the factor $\left(1 - \mathsf{E}\,(r)^2\, h^2\right)$ in the denominator) improved performance for within-population models, but no evidence for the same in across-population models.

- We find evidence that scaling by genetic correlations reduces RMSE slightly in the across-population case.

- We also note that both models have an $R^2 > 0.99$, implying a lack of uncaptured interaction effects between the different choices.

The best-fitting expected accuracies for the within-population models and the across-population models are shown in Figure S4, that is, the expected accuracies obtained using the combination $c$ with the lowest $\delta_c^{\mathcal{M}_{\text{within}}}$ and $\delta_c^{\mathcal{M}_{\text{across}}}$, respectively.[1] Evidently, the expected accuracy formula (S2) can perform well for within-population GP. However, even in the best case scenario, the formulas fail to predict the accuracy of the across-population models, usually by overestimating the accuracy. Note that Figure S4 does not necessarily imply that the formula is useful in the within-population case. The figure contains a post-hoc best fit, so the only implication is that it is *possible* to find some parameters that make formula (S2) work *given that one knows the realized accuracies.* To demonstrate how impactful the above listed choices can be, Figure S4 also contains the worst-fitting expected accuracies.[2] Here, the expected accuracies completely misses the realized accuracies, both in models predicting within and across populations. So, in conclusion, even under reasonable choices the formulas can perform poorly, and we find no combination of choices that makes the formula perform well for across-population GP.

#### Showing that $R^2 = h^2 r^2$ only if $y_i = g_i + \varepsilon_i$

Derivations of expected accuracy formulas frequently rely on the assumption that $R^2 = r^2 h^2$, where $R$ is the phenotypic prediction accuracy $\mathsf{Corr}\,(y_i, \hat{y}_i)$, $r$ is the GP accuracy $\mathsf{Corr}\,(g_i, \hat{g}_i)$ and $h^2 = \frac{\mathsf{Var}(g_i)}{\mathsf{Var}(y_i)}$ is the heritability. Here we show that the assumption $R^2 = r^2 h^2$ only holds in the case where there are no complicating environmental effects in the model.

We first consider the simple case where $R^2 = r^2 h^2$ does hold. Following the notation from the main

[1] The best set of choices for the within-population models was using $M_{e7}$, with $X_{h^2=h^2_M} = 0$, $X_{h^2_{\text{test}}} = 0$ and $X_{r^2h^2\neq 0} = 0$. The best set of choices for the across-population models was using $M_{e7}$, with $X_{h^2=h^2_M} = 1$, $X_{h^2_{\text{test}}} = 1$, $X_{r^2h^2\neq 0} = 0$ and $X_{\text{gc}} = 0$.

[2] The worst set of choices for the within-population models was using $M_{e5}$, with $X_{h^2=h^2_M} = 1$, $X_{h^2_{\text{test}}} = 0$ and $X_{r^2h^2\neq 0} = 0$. The worst set of choices for the across-population models was using $M_{e6}$, with $X_{h^2=h^2_M} = 0$, $X_{h^2_{\text{test}}} = 0$, $X_{r^2h^2\neq 0} = 1$ and $X_{\text{gc}} = 0$.

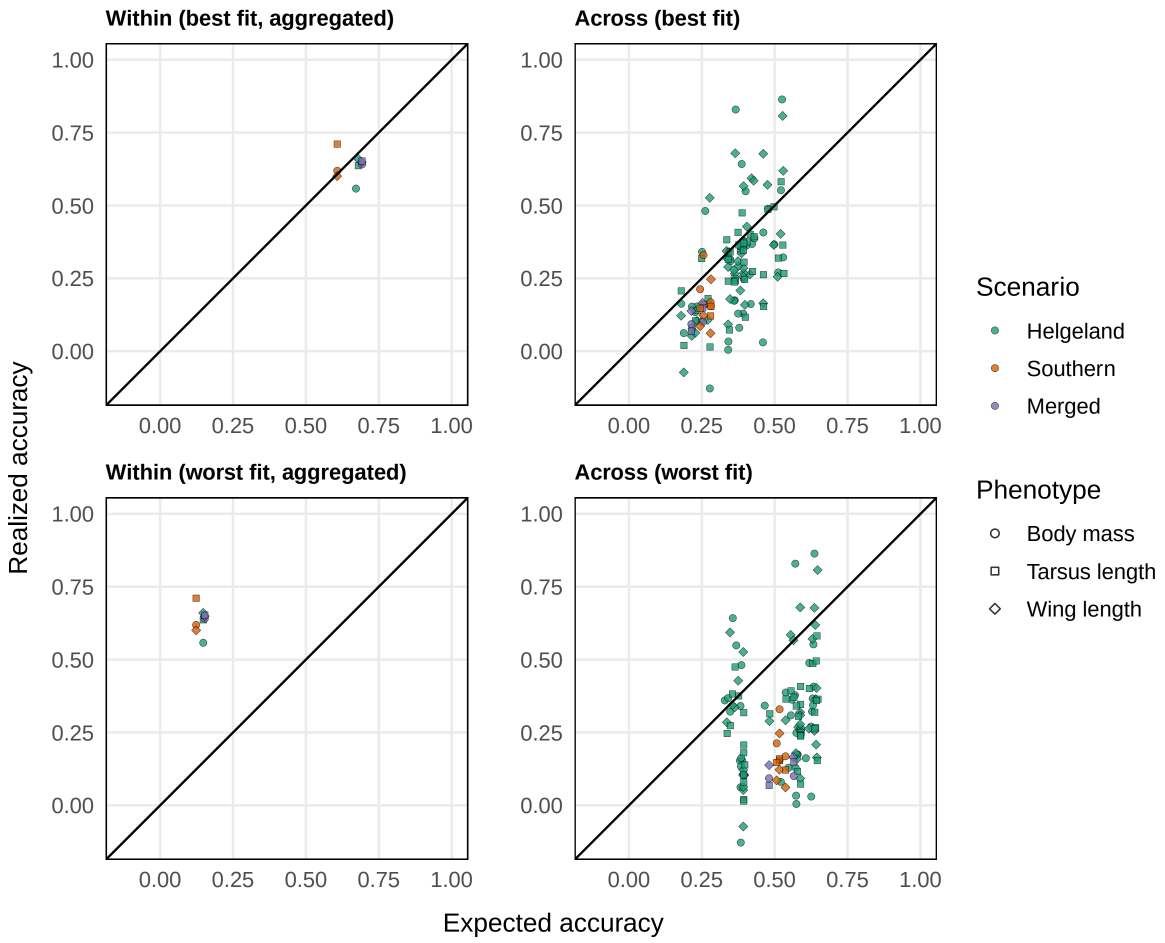


Figure S4: Realized genomic prediction accuracy vs. the expected accuracy given by a version of the expected accuracy formula. The x-axis in the top panels use the expected accuracy formula that gives the best fit (lowest RMSE) to the realized accuracies from the within-population models (left) and the across-population models (right), while the bottom panel uses the parameters that give the worst fit (highest RMSE). In the left panels, the each point corresponds to (expected or realized) accuracies averaged over the 10 folds in a given model, while in the right panels, the points corresponds to (expected or realized) accuracies in single models. Diagonals are shown as black lines, so a perfect fit would follow these lines.

text, we let

$$y_i = g_i + \epsilon_i,$$

*i.e.*, the trait is determined only by the genetic value and a homogeneous environmental effect. Then

$$\hat{y}_i = \hat{g}_i,$$

since the environment tells us nothing about the trait. Then

$$
\begin{aligned}
R^2 = \mathsf{Corr}\left(y_i, \hat{y}_i\right) &= \frac{\mathsf{Cov}\left(g_i + \epsilon_i,\ \hat{g}_i\right)}{\sqrt{\mathsf{Var}\left(y_i\right)\mathsf{Var}\left(\hat{g}_i\right)}} \\
&= \frac{\mathsf{Cov}\left(g_i, \hat{g}_i\right)}{\sqrt{\mathsf{Var}\left(y_i\right)\mathsf{Var}\left(\hat{g}_i\right)}} \\
&= \frac{\mathsf{Cov}\left(g_i, \hat{g}_i\right)}{\sqrt{\mathsf{Var}\left(g_i\right)\mathsf{Var}\left(\hat{g}_i\right)}} \cdot \sqrt{\frac{\mathsf{Var}\left(g_i\right)}{\mathsf{Var}\left(y_i\right)}} \\
&= \mathsf{Corr}\left(g_i, \hat{g}_i\right) \cdot h = r \cdot h \\
\Rightarrow R^2 &= r^2 h^2 \ .
\end{aligned}
$$

Conversely, if

$$y_i = x_i + g_i + \epsilon_i,$$

*i.e.*, $i$ in addition to its genetic value $g_i$ the trait value of $i$ is determined by other (fixed or random) effects, which we lump into a single term $x_i$, then

$$\hat{y}_i = \hat{x}_i + \hat{g}_i.$$

In this case,

$$
\begin{aligned}
\mathsf{Corr}\left(y_i, \hat{y}_i\right) &= \frac{\mathsf{Cov}\left(x_i + g_i + \epsilon_i, \hat{x}_i + \hat{g}_i\right)}{\sqrt{\mathsf{Var}\left(y_i\right)\mathsf{Var}\left(\hat{y}_i\right)}} \\
&= \frac{\mathsf{Cov}\left(x_i, \hat{x}_i\right) + \mathsf{Cov}\left(g_i, \hat{g}_i\right)}{\sqrt{\mathsf{Var}\left(y_i\right)\mathsf{Var}\left(\hat{y}_i\right)}} \\
&= \left(\frac{\mathsf{Cov}\left(x_i, \hat{x}_i\right)}{\sqrt{\mathsf{Var}\left(y_i\right)\mathsf{Var}\left(\hat{y}_i\right)}} \cdot h\right) + \left(\mathsf{Corr}\left(g_i, \hat{g}_i\right) \cdot h \cdot \frac{\mathsf{Var}\left(\hat{g}_i\right)}{\mathsf{Var}\left(\hat{y}_i\right)}\right) \\
&= a \cdot h + r \cdot h \cdot b \\
\Rightarrow R^2 &= \left(a + r \cdot h \cdot b\right)^2 ,
\end{aligned}
$$

where $a = \frac{\mathsf{Cov}(x_i, \hat{x}_i)}{\sqrt{\mathsf{Var}(y_i)\mathsf{Var}(\hat{y}_i)}}$ and $b = \frac{\mathsf{Var}(\hat{g}_i)}{\mathsf{Var}(\hat{y}_i)}$. Therefore, we will only have $R^2 = r^2 h^2$ if $a = 0$ and $b = 1$. For $a = 0$ we must have, $\mathsf{Cov}(x_i, \hat{x}_i) = 0$, so this situation once again reduces to a trivial case where we have no information about $x_i$. To conclude, we will only have $R^2 = h^2 r^2$ in a situation where $y_i = g_i + \epsilon_i$.

### S4: Additional population-differentiation measures

#### Genetic correlations

In the main text we assessed the impact of population-differences in relatedness, LD, and fixation indices on the across-population GP accuracy. Another potential factor in determining said accuracy is population-differences in the causal effects of the SNP markers (Wientjes *et al.*, 2015; Wang *et al.*, 2020). Such differences can be quantified with trait-specific genetic correlations between the populations, which measure whether the focal trait has the same genetic basis in the two populations.

To find the genetic correlations between the populations, we treated the measurements of a given trait from the two populations as two different traits. We estimated the genetic correlations using bivariate genomic animal models with response $\left(y_{ijk}^{(1)},\ y_{ijk}^{(2)}\right)^{\top}$, where the superscript (1) denotes a measurement in the training set and (2) a measurement in the test set. The bivariate animal models included all the same effects as (1), which we modeled jointly in the two populations, except $g_i$ and $\varepsilon_{ijk}$ which we replaced by bivariate effects $\left(g_i^{(1)},\ g_i^{(2)}\right)^{\top}$ and $\left(\varepsilon_{ijk}^{(1)},\ \varepsilon_{ijk}^{(2)}\right)^{\top}$. For these bivariate effects we estimated correlations between the populations, respectively genetic correlations $\rho_g$ and residual (environmental) correlations $\rho_\varepsilon$. The bivariate models were fitted in a Bayesian framework using `R-INLA`, using the reparameterization method given by Mathew *et al.* (2016). In their notation, we used $\mathsf{Inv\text{-}Gamma}(\alpha = 5; \beta = 1.2)$ priors on $\sigma_{a_1}^2$ and $\sigma_{e_1}^2$, $\mathsf{Inv\text{-}Gamma}(\alpha = 5; \beta = 0.2)$ priors on $\sigma_{a_2}^2$ and $\sigma_{e_2}^2$, and a $\mathsf{N}\left(1, 0.3^2\right)$ prior on $\kappa_{12}$. The remaining details of the models are similar to the univariate models which we described in Appendix S1.

These bivariate models come with some caveats. Firstly, for the effects which were not treated as bivariate, it would have been preferable to estimate them independently in the two populations, rather than jointly. However, the former approach leads to a large amount of hyperparameters in `R-INLA`, and thus led to non-convergence for the bivariate models in Scenario 3 (Merged). For the models in Scenario 1 and 2, the results were similar with the two approaches (not shown), so we present the results from the models with joint effects, which always converged. Secondly, a drawback of the reparameterization by Mathew *et al.* (2016) is that the treatment of the two populations is not symmetric. That is, flipping which population is treated as (1) or (2) would result in a different result for the genetic correlations (which is apparent in the results for Scenario 3, see below). Thirdly, as pointed out in the main text, fitting these bivariate models relies on phenotypic measurements from both the training and test populations, so the genetic correlations are not relevant in providing *a priori* estimates of across-population GP accuracy: the phenotypes in the test set could be used to train a GP model, so this would not be in an across-population GP scenario. And, since the genetic correlations are specific to a given trait, estimates from previous across-population GP models with different traits cannot be used to measure the genetic correlation in a new study.

Figure S5 shows the across-population accuracies plotted against the estimated genetic correlations, similarly to Figure 4 in the main text. There seems to be a slightly negative relationship between the accuracy and genetic correlations. This result is contrary to the theoretical expectation, which is that the accuracy would be higher for higher genetic correlations (Wientjes *et al.*, 2015). This seems to imply that the reason that across-population accuracy suffers is *not* due to a difference in genetic bases for the traits between populations, but again we note that the genetic correlation models should be taken with a grain of salt due to the aforementioned issues. Figure S5 also shows that the residual environmental correlations have little impact on the across-population accuracy.

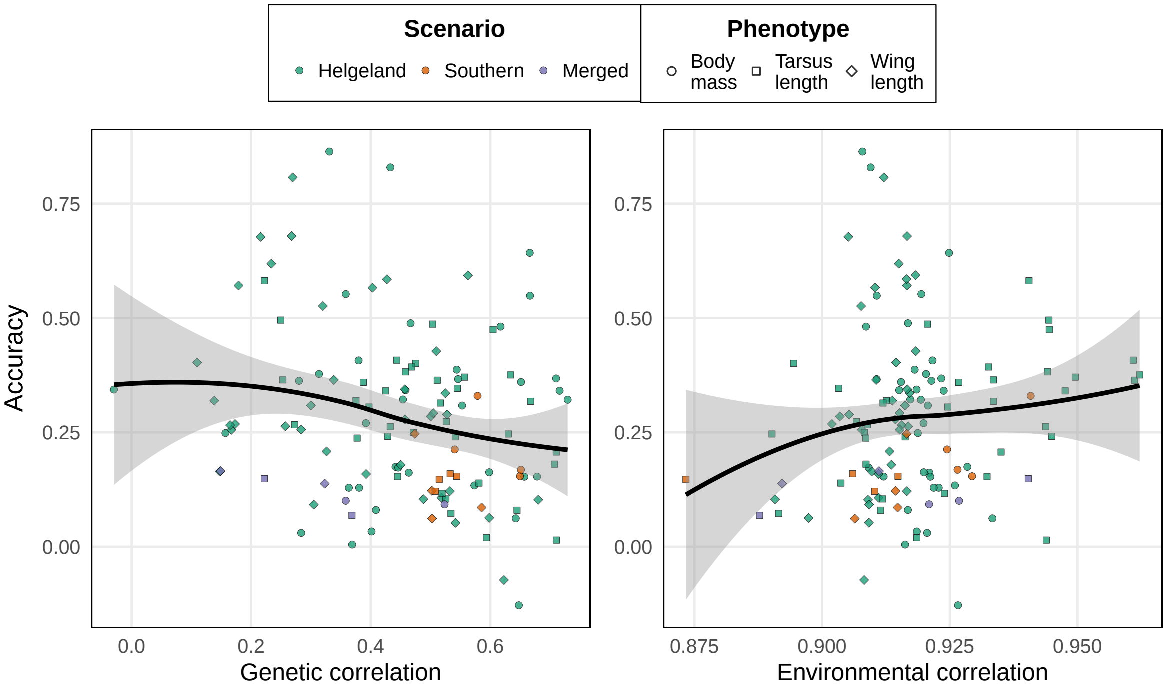


Figure S5: Scatter plot of across-population prediction accuracies vs estimated trait-specific genetic correlations and residual environmental correlations.

#### Heritability ratios

The full genomic animal models from S1 were used to derive heritabilities $h^2_{\mathrm{AM}}$ in the test set that could be compared to the test set heritabilities $h^2_{\mathrm{GP}}$ derived from the corresponding GP model. The ratio $h^2_{\mathrm{GP}}/h^2_{\mathrm{AM}}$ then indicates to what degree the GP model captured the heritability in the test set (where phenotypes per definition are unknown), compared to the situation where the test set individuals also had known phenotypes (*i.e.*, the full genomic animal models in S1). As seen in Figure S6, the ratio $h^2_{\mathrm{GP}}/h^2_{\mathrm{AM}}$ seemed to relate to across-population GP accuracy. The best-performing models had a value of $h^2_{\mathrm{GP}}/h^2_{\mathrm{AM}}$ close to 1, while we got low-to-moderate accuracies when the heritability was underestimated ($h^2_{\mathrm{GP}}/h^2_{\mathrm{AM}} < 1$), and moderate accuracies when the heritability was overestimated ($h^2_{\mathrm{GP}}/h^2_{\mathrm{AM}} > 1$).

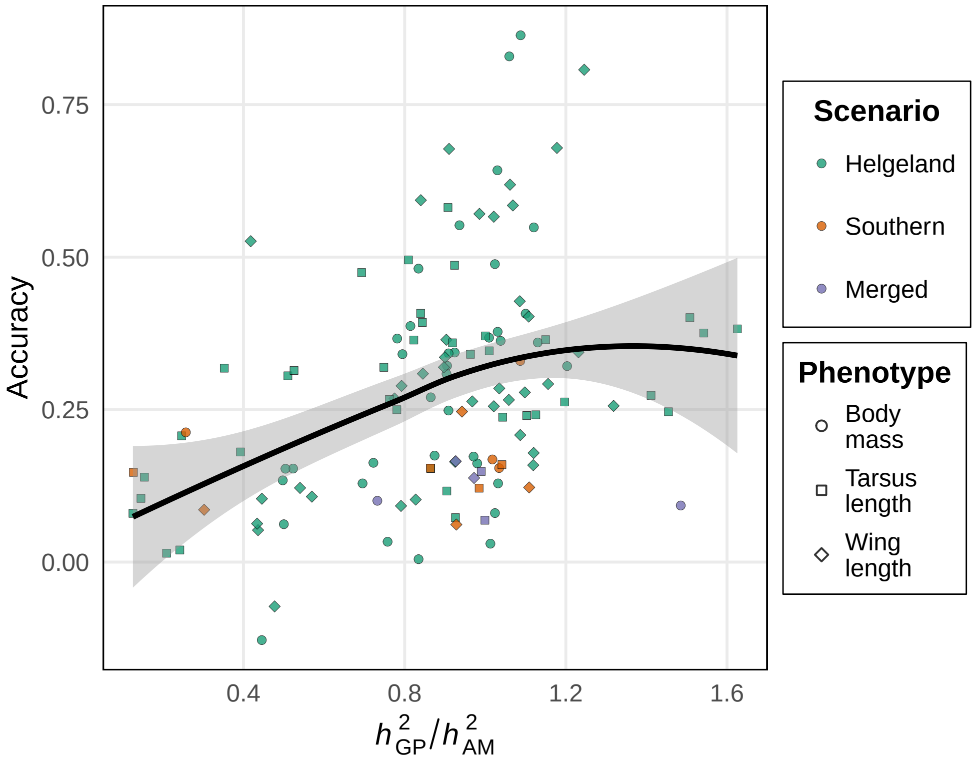


Figure S6: Scatter plot of across-population prediction accuracies vs the ratio $h^2_{\mathrm{GP}}/h^2_{\mathrm{AM}}$ between the test set heritability captured by the GP models and an animal models with access to the test set phenotypes.

### S5: Variable importance for across-population accuracy

In the main text we investigated how various parameters impacted the across-population genomic prediction accuracy, namely the number of individuals in the training set $N$, $\overline{G}_{\mathrm{ac}}$, $\tilde{G}_{\mathrm{ac}}$, $\delta$LD, $\Delta$LD, and $F_{st}$, while in Appendix S4 we also investigated the impact of genetic correlations $\rho_g$ and heritability ratios $h^2_{\mathrm{GP}}/h^2_{\mathrm{AM}}$. Figure S7 shows a pairs plot between all these aforementioned parameters, expected accuracy $\mathsf{E}(r)$ and realized accuracy $r$, with scatter plots in the lower triangular part, pairwise sample correlations on the upper diagonal and histograms of each variable on the diagonal. In this calculation, we used best-performing estimator for $\mathsf{E}(r)$ in the analysis in Appendix S3. Beyond what was discussed in the main text, we can see that $F_{st}$ and $\delta$LD are highly correlated, while $\Delta$LD and $\mathsf{E}(r)$ are highly negative correlated, with a higher absolute correlation than either of the two parameters have with realized accuracies $r$. Thus, $\delta$LD seems to mostly reflect allele-frequency differences, while $\Delta$LD contains much of the same information as is encoded in $\mathsf{E}(r)$.

To see how the various parameters impact the accuracy, we fitted several multiple regression models with the across-population accuracy as response. We included four different sets of covariates in different models, and for each set we fit both a linear model (LM), as well as a generalized additive model (GAM) to capture potential non-linearity in the effects of the covariates. In *Model 1*, we included as covariates the parameters used in the main text, namely $N$, $\overline{G}_{\text{ac}}$, $\tilde{G}_{\text{ac}}$, $\delta$LD, $\Delta$LD, and $F_{st}$. In *Model 2* we included the same covariates, in addition to parameter depending on phenotype data, namely genetic correlation $\rho_g$, heritability in the test set $h^2_{\text{AM}}$ (derived from the full animal models in Appendix S1), and the heritability ratio $h^2_{\text{GP}}/h^2_{\text{AM}}$. *Model 3* contained only the expected accuracy $\mathsf{E}(r)$, again using the best performing choices from Appendix S3, and was only fitted as a linear model since $\mathsf{E}(r)$ should directly predict $r$. *Model 4* contained all previously mentioned covariates, including $\mathsf{E}(r)$, and excluding the parameters used to compute $\mathsf{E}(r)$, namely $N$, $h^2_{\text{AM}}$ and $\tilde{G}_{\text{ac}}$. The GAM for *Model 4* imposed a linear effect for $\mathsf{E}(r)$ for the same reason that we did not fit a GAM for *Model 3*. We used hierarchical partitioning, as implemented in the `R` packages `glmm.hp` (Lai *et al.*, 2022) and `gam.hp` (Lai *et al.*, 2024), to partition the variance and deviance explained by the LMs and GAMs, respectively, into parts attributable to each covariate.

Table S3 contains the results from the variable importance analysis, that is, from the hierarchical partitioning of variance (LMs) or deviance (GAMs) explained by Models 1 through 4. We found that $\overline{G}_{\text{ac}}$, $\tilde{G}_{\text{ac}}$, and $\Delta$LD were the most important variables in all LM where they were included. $\delta$LD and $F_{st}$ were only given importance in the non-linear models. Notably, the LMs never explain more than 40% of the variance, and even when using GAMs, we at best only explain about 55% of the deviance (Model 4). The additional information from consider the parameters relying on phenotypes was marginal, as found by comparing models 1 and 2. Comparing Models 1 and 3, we found that our parameters together explain more variance in the across-population than the formula for expected accuracy does.

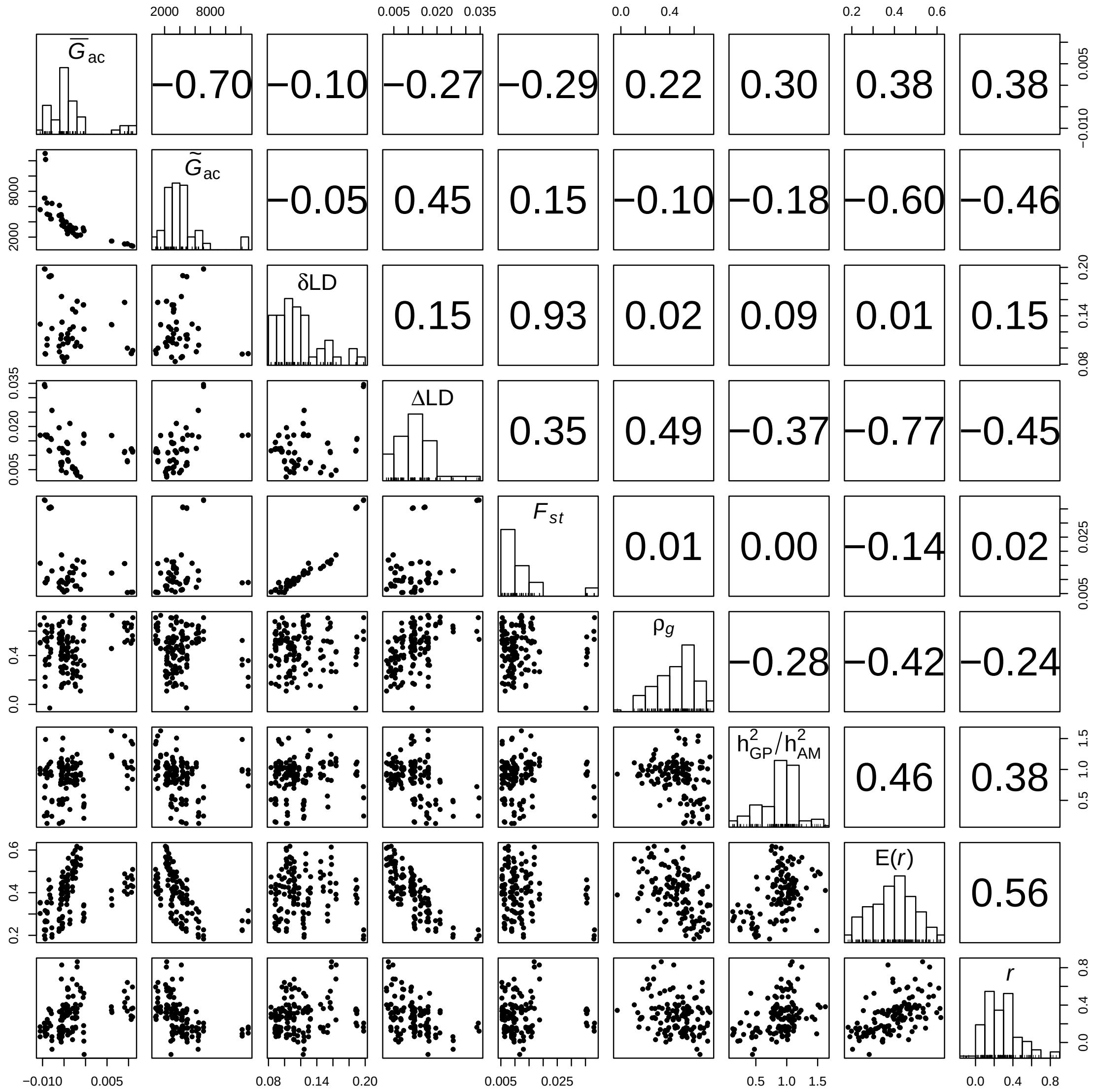


Figure S7: Pairwise scatter plots (lower triangular matrix) and sample Pearson correlations (upper triangular matrix) between population-differentiation measures ($\overline{G}_{\mathrm{ac}}$, $\tilde{G}_{\mathrm{ac}}$, $\delta$LD, $\Delta$LD, $F_{st}$, genetic correlation $\rho_g$, heritability ratio $h^2_{\mathrm{GP}}/h^2_{\mathrm{AM}}$), expected accuracy $\mathsf{E}(r)$ and realized accuracy $r$ in the across-population models. The diagonals show histograms for each variable.

Table S3: Results from a variable importance analysis of the impact of various parameters upon across-population GP prediction accuracy. Each row shows how much variance or deviance is explained by a given covariate in a given model for the LMs or GAMs, respectively, according to hierarchical partitioning. The percentages in parentheses indicates what proportion of the total variance/deviance explained by that model is attributed to the given covariate. A dash indicates the covariate was not included in the given model. The bottom row indicates how much variance/deviance was explained by the LMs/GAMs, respectively.

| | Model 1 | | Model 2 | | Model 3 | Model 4 | |
|---|---|---|---|---|---|---|---|
| | Linear | GAM | Linear | GAM | Linear | Linear | GAM |
| $N$ | 0.0582 (15.54%) | 0.0634 (12.12%) | 0.0330 (8.42%) | 0.0423 (7.77%) | - | - | - |
| $\overline{G}_{\mathrm{ac}}$ | 0.0884 (23.61%) | 0.0638 (12.20%) | 0.0751 (19.16%) | 0.0595 (10.93%) | - | 0.0803 (21.37%) | 0.0815 (14.78%) |
| $\tilde{G}_{\mathrm{ac}}$ | 0.0979 (26.15%) | 0.0987 (18.87%) | 0.0972 (24.80%) | 0.0910 (16.72%) | - | - | - |
| $\delta$LD | 0.0238 (6.36%) | 0.1086 (20.76%) | 0.0217 (5.54%) | 0.1050 (19.29%) | - | 0.0259 (6.89%) | 0.1244 (22.56%) |
| $\Delta$LD | 0.0915 (24.44%) | 0.1009 (19.29%) | 0.0707 (18.04%) | 0.0859 (15.78%) | - | 0.0735 (19.56%) | 0.1010 (18.32%) |
| $F_{st}$ | 0.0146 (3.90%) | 0.0876 (16.75%) | 0.0127 (3.24%) | 0.0884 (16.24%) | - | 0.0146 (3.89%) | 0.1072 (19.44%) |
| $\rho_g$ | - | - | 0.0268 (6.84%) | 0.0211 (3.88%) | - | 0.0256 (6.81%) | 0.0307 (5.57%) |
| $h^2_{\mathrm{AM}}$ | - | - | 0.0013 (0.33%) | 0.0092 (1.69%) | - | - | - |
| $h^2_{\mathrm{GP}}/h^2_{\mathrm{AM}}$ | - | - | 0.0535 (13.65%) | 0.0419 (7.70%) | - | 0.0524 (13.94%) | 0.0495 (8.98%) |
| $\mathsf{E}(r)$ | - | - | - | - | 0.2620 (100%) | 0.1035 (27.54%) | 0.0570 (10.34%) |
| Total | 0.3744 (100%) | 0.5230 (100%) | 0.3921 (100%) | 0.5443 (100%) | 0.2620 (100%) | 0.3758 (100%) | 0.5513 (100%) |

### S6: Supplemental tables

#### Genomic animal models

##### Body mass

Table S4: Statistics related to the genomic animal models with body mass as response. For statistics derived from the model, we show posterior means and the 0.025 and 0.975 quantiles of the posterior distribution in parenthesis. $N$ is the number of individuals, $N_{\text{obs}}$ is the number of observations, $M$ is the number of SNPs used, $d$ is the value that was added to the diagonal of the GRM for the respective model, $\mu$ is the intercept, $\beta_{\text{sex=m}}$ is the effect of being male compared to female, $\beta_{\text{monthX}}$ is the effect of month $X$ compared to January, $\beta_{\text{age}}$ is the effect of age, $\sigma^2_{\text{G}}$ is the additive genetic variance, $\sigma^2_{\text{hatch-year}}$ is the variance explained by hatch year, $\sigma^2_{\text{island}}$ is the variance explained by island of measurement, $\sigma^2_{\text{id}}$ is the variance explained by permanent environmental effects, $\sigma^2_{\text{session}}$ is the variance explained by measurement session and $\sigma^2_{\varepsilon}$ is the residual variance.

| Scenario | Helgeland | Southern | Merged |
|---|---|---|---|
| $N$ | 3456 | 2254 | 5710 |
| $N_{\text{obs}}$ | 7977 | 4124 | 12101 |
| $M$ | 65221 | 64917 | 64943 |
| $d$ | 6.6e-03 | 5.0e-03 | 6.2e-03 |
| $\mu$ | 32.22 (30.96, 33.47) | 30.51 (29.83, 31.19) | 31.41 (30.74, 32.09) |
| $\beta_{\text{sex=m}}$ | -0.32 (-0.44, -0.20) | 0.41 (0.28, 0.54) | -0.06 (-0.16, 0.03) |
| $\beta_{\text{month=2}}$ | 0.12 (-1.27, 1.52) | 0.11 (-0.53, 0.75) | -0.09 (-0.75, 0.57) |
| $\beta_{\text{month=3}}$ | -0.33 (-1.98, 1.33) | -0.35 (-1.00, 0.30) | -0.57 (-1.24, 0.10) |
| $\beta_{\text{month=4}}$ | -0.39 (-1.72, 0.93) | -0.25 (-1.03, 0.54) | -0.13 (-0.87, 0.61) |
| $\beta_{\text{month=5}}$ | 0.10 (-1.14, 1.34) | 1.34 (0.41, 2.27) | 0.63 (-0.05, 1.30) |
| $\beta_{\text{month=6}}$ | -0.28 (-1.52, 0.96) | 1.22 (0.32, 2.13) | 0.26 (-0.42, 0.93) |
| $\beta_{\text{month=7}}$ | -0.80 (-2.04, 0.43) | -0.57 (-1.42, 0.29) | -0.29 (-0.96, 0.39) |
| $\beta_{\text{month=8}}$ | -0.37 (-1.61, 0.87) | -0.09 (-1.25, 1.07) | 0.16 (-0.52, 0.83) |
| $\beta_{\text{month=9}}$ | 0.23 (-1.02, 1.49) | 1.62 (0.76, 2.48) | 0.81 (0.12, 1.51) |
| $\beta_{\text{month=10}}$ | 0.10 (-1.14, 1.34) | 1.51 (0.69, 2.33) | 0.63 (-0.05, 1.31) |
| $\beta_{\text{month=11}}$ | 0.12 (-1.18, 1.41) | 0.89 (0.03, 1.76) | 0.63 (-0.10, 1.37) |
| $\beta_{\text{age}}$ | 0.05 (0.01, 0.09) | 0.09 (0.05, 0.13) | 0.05 (0.02, 0.08) |
| $\sigma^2_{\text{G}}$ | 1.30 (1.07, 1.58) | 1.42 (1.18, 1.70) | 1.24 (1.07, 1.43) |
| $\sigma^2_{\text{hatch-year}}$ | 0.09 (0.04, 0.18) | 0.07 (0.01, 0.24) | 0.06 (0.02, 0.12) |
| $\sigma^2_{\text{island}}$ | 0.07 (0.01, 0.21) | 0.02 (0.00, 0.10) | 0.05 (0.01, 0.15) |
| $\sigma^2_{\text{id}}$ | 0.71 (0.53, 0.92) | 0.35 (0.20, 0.56) | 0.66 (0.53, 0.80) |
| $\sigma^2_{\text{session}}$ | 2.72 (2.59, 2.85) | 1.32 (1.21, 1.44) | 2.30 (2.21, 2.40) |
| $\sigma^2_{\varepsilon}$ | 0.21 (0.19, 0.23) | 0.28 (0.24, 0.32) | 0.23 (0.21, 0.25) |

#### Tarsus length

Table S5: Statistics related to the genomic animal models with tarsus length as response. For statistics derived from the model, we show posterior means and the 0.025 and 0.975 quantiles of the posterior distribution in parenthesis. $N$ is the number of individuals, $N_{\text{obs}}$ is the number of observations, $M$ is the number of SNPs used, $d$ is the value that was added to the diagonal of the GRM for the respective model, $\mu$ is the intercept, $\beta_{\text{sex=m}}$ is the effect of being male compared to female, $\beta_{\text{monthX}}$ is the effect of month $X$ compared to January, $\beta_{\text{age}}$ is the effect of age, $\sigma^2_{\text{G}}$ is the additive genetic variance, $\sigma^2_{\text{hatch-year}}$ is the variance explained by hatch year, $\sigma^2_{\text{island}}$ is the variance explained by island of measurement, $\sigma^2_{\text{id}}$ is the variance explained by permanent environmental effects, $\sigma^2_{\text{session}}$ is the variance explained by measurement session and $\sigma^2_{\varepsilon}$ is the residual variance.

| Scenario | Helgeland | Southern | Merged |
|---|---|---|---|
| $N$ | 3445 | 2255 | 5700 |
| $N_{\text{obs}}$ | 8067 | 4171 | 12238 |
| $M$ | 65221 | 64917 | 64943 |
| $d$ | 6.6e-03 | 5.0e-03 | 6.1e-03 |
| $\mu$ | 19.23 (19.07, 19.38) | 19.46 (19.32, 19.61) | 19.35 (19.26, 19.45) |
| $\beta_{\text{sex=m}}$ | 0.08 (0.03, 0.13) | 0.11 (0.05, 0.17) | 0.09 (0.06, 0.13) |
| $\beta_{\text{month=2}}$ | 0.01 (-0.31, 0.34) | 0.03 (-0.07, 0.13) | 0.07 (-0.01, 0.16) |
| $\beta_{\text{month=3}}$ | 0.32 (0.14, 0.51) | -0.02 (-0.12, 0.08) | 0.02 (-0.06, 0.11) |
| $\beta_{\text{month=4}}$ | 0.12 (-0.03, 0.28) | -0.03 (-0.17, 0.10) | 0.05 (-0.05, 0.14) |
| $\beta_{\text{month=5}}$ | 0.14 (-0.00, 0.28) | 0.19 (0.05, 0.33) | 0.08 (-0.01, 0.16) |
| $\beta_{\text{month=6}}$ | 0.19 (0.05, 0.34) | 0.25 (0.11, 0.39) | 0.13 (0.04, 0.21) |
| $\beta_{\text{month=7}}$ | 0.21 (0.07, 0.36) | 0.18 (0.05, 0.31) | 0.15 (0.06, 0.23) |
| $\beta_{\text{month=8}}$ | 0.22 (0.08, 0.36) | 0.09 (-0.08, 0.26) | 0.15 (0.06, 0.23) |
| $\beta_{\text{month=9}}$ | 0.32 (0.18, 0.46) | -0.01 (-0.14, 0.12) | 0.23 (0.15, 0.32) |
| $\beta_{\text{month=10}}$ | 0.26 (0.12, 0.40) | 0.13 (0.00, 0.25) | 0.19 (0.11, 0.28) |
| $\beta_{\text{month=11}}$ | 0.19 (0.04, 0.34) | 0.09 (-0.04, 0.22) | 0.13 (0.04, 0.22) |
| $\beta_{\text{age}}$ | 0.00 (-0.00, 0.01) | -0.02 (-0.02, -0.01) | -0.01 (-0.01, -0.00) |
| $\sigma^2_{\text{G}}$ | 0.25 (0.21, 0.29) | 0.24 (0.20, 0.30) | 0.24 (0.21, 0.27) |
| $\sigma^2_{\text{hatch-year}}$ | 0.01 (0.00, 0.02) | 0.02 (0.00, 0.06) | 0.01 (0.00, 0.02) |
| $\sigma^2_{\text{island}}$ | 0.00 (0.00, 0.01) | 0.00 (0.00, 0.02) | 0.00 (0.00, 0.01) |
| $\sigma^2_{\text{id}}$ | 0.35 (0.33, 0.39) | 0.34 (0.30, 0.38) | 0.35 (0.33, 0.38) |
| $\sigma^2_{\text{session}}$ | 0.02 (0.02, 0.02) | 0.02 (0.01, 0.02) | 0.02 (0.02, 0.02) |
| $\sigma^2_{\varepsilon}$ | 0.01 (0.01, 0.01) | 0.02 (0.01, 0.02) | 0.01 (0.01, 0.02) |

#### Wing length

Table S6: Statistics related to the genomic animal models with wing length as response. For statistics derived from the model, we show posterior means and the 0.025 and 0.975 quantiles of the posterior distribution in parenthesis. $N$ is the number of individuals, $N_{\text{obs}}$ is the number of observations, $M$ is the number of SNPs used, $d$ is the value that was added to the diagonal of the GRM for the respective model, $\mu$ is the intercept, $\beta_{\text{sex=m}}$ is the effect of being male compared to female, $\beta_{\text{monthX}}$ is the effect of month $X$ compared to January, $\beta_{\text{age}}$ is the effect of age, $\sigma^2_{\text{G}}$ is the additive genetic variance, $\sigma^2_{\text{hatch-year}}$ is the variance explained by hatch year, $\sigma^2_{\text{island}}$ is the variance explained by island of measurement, $\sigma^2_{\text{id}}$ is the variance explained by permanent environmental effects, $\sigma^2_{\text{session}}$ is the variance explained by measurement session and $\sigma^2_{\varepsilon}$ is the residual variance.

| Scenario | Helgeland | Southern | Merged |
|---|---|---|---|
| $N$ | 3424 | 2254 | 5678 |
| $N_{\text{obs}}$ | 7997 | 4151 | 12148 |
| $M$ | 65220 | 64917 | 64943 |
| $d$ | 6.6e-03 | 5.0e-03 | 6.1e-03 |
| $\mu$ | 78.00 (77.08, 78.91) | 77.66 (77.07, 78.25) | 77.63 (77.10, 78.16) |
| $\beta_{\text{sex=m}}$ | 2.73 (2.64, 2.83) | 2.67 (2.55, 2.78) | 2.71 (2.63, 2.78) |
| $\beta_{\text{month=2}}$ | 0.77 (-0.94, 2.48) | 0.78 (0.24, 1.31) | 0.64 (0.15, 1.13) |
| $\beta_{\text{month=3}}$ | -0.20 (-1.34, 0.94) | 0.71 (0.16, 1.25) | 0.53 (0.03, 1.04) |
| $\beta_{\text{month=4}}$ | -0.33 (-1.28, 0.61) | 0.87 (0.21, 1.54) | 0.28 (-0.28, 0.83) |
| $\beta_{\text{month=5}}$ | 0.07 (-0.81, 0.95) | 0.71 (-0.06, 1.47) | 0.41 (-0.09, 0.92) |
| $\beta_{\text{month=6}}$ | -0.19 (-1.08, 0.69) | 1.07 (0.31, 1.84) | 0.16 (-0.34, 0.66) |
| $\beta_{\text{month=7}}$ | -0.42 (-1.30, 0.47) | -0.75 (-1.46, -0.04) | -0.08 (-0.59, 0.42) |
| $\beta_{\text{month=8}}$ | -0.35 (-1.23, 0.54) | -1.12 (-2.04, -0.19) | -0.01 (-0.52, 0.49) |
| $\beta_{\text{month=9}}$ | -1.11 (-2.01, -0.22) | 0.10 (-0.70, 0.90) | -0.72 (-1.24, -0.20) |
| $\beta_{\text{month=10}}$ | 0.25 (-0.64, 1.13) | 0.16 (-0.52, 0.84) | 0.58 (0.07, 1.08) |
| $\beta_{\text{month=11}}$ | 0.65 (-0.27, 1.57) | 0.74 (0.02, 1.46) | 0.95 (0.40, 1.50) |
| $\beta_{\text{age}}$ | 0.33 (0.30, 0.35) | 0.26 (0.22, 0.29) | 0.31 (0.29, 0.33) |
| $\sigma^2_{\text{G}}$ | 1.50 (1.32, 1.72) | 1.26 (1.06, 1.50) | 1.32 (1.18, 1.47) |
| $\sigma^2_{\text{hatch-year}}$ | 0.05 (0.02, 0.12) | 0.12 (0.04, 0.28) | 0.11 (0.05, 0.22) |
| $\sigma^2_{\text{island}}$ | 0.14 (0.04, 0.37) | 0.03 (0.00, 0.19) | 0.09 (0.03, 0.21) |
| $\sigma^2_{\text{id}}$ | 0.30 (0.20, 0.42) | 0.25 (0.13, 0.41) | 0.33 (0.25, 0.43) |
| $\sigma^2_{\text{session}}$ | 0.92 (0.84, 0.99) | 0.70 (0.61, 0.79) | 0.86 (0.80, 0.92) |
| $\sigma^2_{\varepsilon}$ | 0.49 (0.45, 0.54) | 0.36 (0.31, 0.42) | 0.45 (0.42, 0.49) |

#### Within-population GP: Body mass

Table S7: Statistics related to each within-population GP model with body mass as response. $N$ is the number of individuals in the training set, $N_{\text{test}}$ is the number of individuals in the test set, $N_{\text{obs}}$ is the number of observations in the training set, $M$ is the number of SNPs used, $\hat{r}$ is the estimated GP accuracy, $\hat{\lambda}$ is the scaling factor used in finding $\hat{r}$ as described in the main text, $\overline{G}_{\text{ac}}$ is the mean relatedness between the training and test set individuals $\tilde{G}_{\text{ac}}$ is the precision of the relatedness between training and test set individuals and $d$ is the value that was added to the diagonal of the GRM for the respective model.

| Scenario | $N$ | $N_{\text{test}}$ | $N_{\text{obs}}$ | $M$ | $\hat{r}$ | $\hat{\lambda}$ | $\overline{G}_{\text{ac}}$ | $\tilde{G}_{\text{ac}}$ | $d$ |
|---|---|---|---|---|---|---|---|---|---|
| Helgeland | 3110 | 346 | 7189 | 65221 | 0.68 | 0.51 | -3.0E-04 | 994.44 | 6.6E-03 |
| Helgeland | 3110 | 346 | 7183 | 65221 | 0.51 | 0.46 | -2.9E-04 | 1025.52 | 6.6E-03 |
| Helgeland | 3111 | 345 | 7183 | 65221 | 0.52 | 0.52 | -3.1E-04 | 967.35 | 6.6E-03 |
| Helgeland | 3110 | 346 | 7118 | 65221 | 0.58 | 0.52 | -2.9E-04 | 969.79 | 6.6E-03 |
| Helgeland | 3111 | 345 | 7193 | 65221 | 0.63 | 0.49 | -3.0E-04 | 973.11 | 6.6E-03 |
| Helgeland | 3110 | 346 | 7148 | 65221 | 0.57 | 0.51 | -3.0E-04 | 973.66 | 6.6E-03 |
| Helgeland | 3111 | 345 | 7219 | 65221 | 0.55 | 0.49 | -2.9E-04 | 927.85 | 6.6E-03 |
| Helgeland | 3110 | 346 | 7166 | 65221 | 0.38 | 0.55 | -3.2E-04 | 1049.53 | 6.6E-03 |
| Helgeland | 3111 | 345 | 7154 | 65221 | 0.46 | 0.54 | -3.0E-04 | 985.07 | 6.6E-03 |
| Helgeland | 3110 | 346 | 7240 | 65221 | 0.68 | 0.50 | -2.8E-04 | 977.08 | 6.6E-03 |
| Southern | 2028 | 226 | 3743 | 64917 | 0.47 | 0.65 | -4.6E-04 | 957.85 | 5.0E-03 |
| Southern | 2029 | 225 | 3705 | 64917 | 0.62 | 0.60 | -4.4E-04 | 949.37 | 5.0E-03 |
| Southern | 2029 | 225 | 3677 | 64917 | 0.52 | 0.64 | -4.4E-04 | 960.83 | 5.0E-03 |
| Southern | 2028 | 226 | 3705 | 64917 | 0.72 | 0.63 | -4.7E-04 | 940.40 | 5.0E-03 |
| Southern | 2029 | 225 | 3724 | 64917 | 0.50 | 0.67 | -4.5E-04 | 912.83 | 5.0E-03 |
| Southern | 2029 | 225 | 3725 | 64917 | 0.69 | 0.61 | -4.3E-04 | 928.00 | 5.0E-03 |
| Southern | 2028 | 226 | 3695 | 64917 | 0.79 | 0.63 | -4.5E-04 | 922.91 | 5.0E-03 |
| Southern | 2029 | 225 | 3745 | 64917 | 0.68 | 0.59 | -4.6E-04 | 974.15 | 5.0E-03 |
| Southern | 2029 | 225 | 3677 | 64917 | 0.61 | 0.64 | -4.5E-04 | 914.58 | 5.0E-03 |
| Southern | 2028 | 226 | 3720 | 64917 | 0.59 | 0.66 | -4.4E-04 | 946.65 | 5.0E-03 |
| Merged | 5139 | 571 | 10911 | 64943 | 0.66 | 0.51 | -2.0E-04 | 1499.93 | 6.2E-03 |
| Merged | 5139 | 571 | 10914 | 64943 | 0.58 | 0.47 | -1.7E-04 | 1489.37 | 6.2E-03 |
| Merged | 5139 | 571 | 10898 | 64943 | 0.66 | 0.54 | -1.8E-04 | 1499.87 | 6.2E-03 |
| Merged | 5139 | 571 | 10910 | 64943 | 0.68 | 0.50 | -1.8E-04 | 1470.63 | 6.2E-03 |
| Merged | 5139 | 571 | 10890 | 64943 | 0.68 | 0.51 | -1.9E-04 | 1584.31 | 6.2E-03 |
| Merged | 5139 | 571 | 10881 | 64943 | 0.58 | 0.50 | -1.8E-04 | 1543.50 | 6.2E-03 |
| Merged | 5139 | 571 | 10950 | 64943 | 0.62 | 0.54 | -1.8E-04 | 1529.31 | 6.2E-03 |
| Merged | 5139 | 571 | 10803 | 64943 | 0.64 | 0.51 | -1.8E-04 | 1512.03 | 6.2E-03 |
| Merged | 5139 | 571 | 10862 | 64943 | 0.57 | 0.55 | -1.9E-04 | 1463.24 | 6.2E-03 |
| Merged | 5139 | 571 | 10890 | 64943 | 0.73 | 0.52 | -1.8E-04 | 1546.83 | 6.2E-03 |

#### Within-population GP: Tarsus length

Table S8: Statistics related to each within-population GP model with tarsus length as response. $N$ is the number of individuals in the training set, $N_{\text{test}}$ is the number of individuals in the test set, $N_{\text{obs}}$ is the number of observations in the training set, $M$ is the number of SNPs used, $\hat{r}$ is the estimated GP accuracy, $\hat{\lambda}$ is the scaling factor used in finding $\hat{r}$ as described in the main text, $\overline{G}_{\text{ac}}$ is the mean relatedness between the training and test set individuals $\tilde{G}_{\text{ac}}$ is the precision of the relatedness between training and test set individuals and $d$ is the value that was added to the diagonal of the GRM for the respective model.

| Scenario | $N$ | $N_{\text{test}}$ | $N_{\text{obs}}$ | $M$ | $\hat{r}$ | $\hat{\lambda}$ | $\overline{G}_{\text{ac}}$ | $\tilde{G}_{\text{ac}}$ | $d$ |
|---|---|---|---|---|---|---|---|---|---|
| Helgeland | 3100 | 345 | 7213 | 65221 | 0.71 | 0.65 | -2.8E-04 | 952.73 | 6.6E-03 |
| Helgeland | 3101 | 344 | 7276 | 65221 | 0.62 | 0.67 | -3.0E-04 | 966.56 | 6.6E-03 |
| Helgeland | 3100 | 345 | 7282 | 65221 | 0.57 | 0.57 | -3.0E-04 | 1000.38 | 6.6E-03 |
| Helgeland | 3101 | 344 | 7271 | 65221 | 0.71 | 0.56 | -2.8E-04 | 950.44 | 6.6E-03 |
| Helgeland | 3100 | 345 | 7247 | 65221 | 0.64 | 0.64 | -3.1E-04 | 941.31 | 6.6E-03 |
| Helgeland | 3101 | 344 | 7285 | 65221 | 0.63 | 0.66 | -3.0E-04 | 989.68 | 6.6E-03 |
| Helgeland | 3101 | 344 | 7251 | 65221 | 0.56 | 0.68 | -3.1E-04 | 995.56 | 6.6E-03 |
| Helgeland | 3100 | 345 | 7258 | 65221 | 0.61 | 0.59 | -2.9E-04 | 1014.26 | 6.6E-03 |
| Helgeland | 3101 | 344 | 7286 | 65221 | 0.71 | 0.63 | -2.9E-04 | 975.53 | 6.6E-03 |
| Helgeland | 3100 | 345 | 7234 | 65221 | 0.59 | 0.67 | -3.2E-04 | 1010.86 | 6.6E-03 |
| Southern | 2029 | 226 | 3777 | 64917 | 0.61 | 0.65 | -4.3E-04 | 992.45 | 5.0E-03 |
| Southern | 2030 | 225 | 3771 | 64917 | 0.67 | 0.64 | -4.4E-04 | 932.69 | 5.0E-03 |
| Southern | 2029 | 226 | 3729 | 64917 | 0.83 | 0.61 | -4.6E-04 | 954.08 | 5.0E-03 |
| Southern | 2030 | 225 | 3763 | 64917 | 0.89 | 0.60 | -4.4E-04 | 867.55 | 5.0E-03 |
| Southern | 2029 | 226 | 3777 | 64917 | 0.52 | 0.72 | -4.4E-04 | 936.92 | 5.0E-03 |
| Southern | 2030 | 225 | 3765 | 64917 | 0.74 | 0.66 | -4.6E-04 | 936.71 | 5.0E-03 |
| Southern | 2030 | 225 | 3742 | 64917 | 0.68 | 0.69 | -4.4E-04 | 944.22 | 5.0E-03 |
| Southern | 2029 | 226 | 3740 | 64917 | 0.59 | 0.72 | -4.7E-04 | 941.27 | 5.0E-03 |
| Southern | 2030 | 225 | 3746 | 64917 | 0.77 | 0.62 | -4.9E-04 | 948.05 | 5.0E-03 |
| Southern | 2029 | 226 | 3729 | 64917 | 0.80 | 0.66 | -4.2E-04 | 965.73 | 5.0E-03 |
| Merged | 5130 | 570 | 11004 | 64943 | 0.56 | 0.64 | -1.9E-04 | 1505.09 | 6.1E-03 |
| Merged | 5130 | 570 | 11013 | 64943 | 0.71 | 0.66 | -1.8E-04 | 1478.44 | 6.1E-03 |
| Merged | 5130 | 570 | 10996 | 64943 | 0.58 | 0.64 | -1.9E-04 | 1521.87 | 6.1E-03 |
| Merged | 5130 | 570 | 10996 | 64943 | 0.72 | 0.62 | -1.8E-04 | 1492.34 | 6.1E-03 |
| Merged | 5130 | 570 | 11019 | 64943 | 0.70 | 0.60 | -1.8E-04 | 1516.68 | 6.1E-03 |
| Merged | 5130 | 570 | 11017 | 64943 | 0.61 | 0.66 | -1.8E-04 | 1517.12 | 6.1E-03 |
| Merged | 5130 | 570 | 11029 | 64943 | 0.70 | 0.68 | -1.8E-04 | 1500.58 | 6.1E-03 |
| Merged | 5130 | 570 | 11003 | 64943 | 0.70 | 0.64 | -1.8E-04 | 1534.69 | 6.1E-03 |
| Merged | 5130 | 570 | 11021 | 64943 | 0.56 | 0.64 | -1.8E-04 | 1537.99 | 6.1E-03 |
| Merged | 5130 | 570 | 11044 | 64943 | 0.69 | 0.65 | -1.8E-04 | 1486.97 | 6.1E-03 |

#### Within-population GP: Wing length

Table S9: Statistics related to each within-population GP model with wing length as response. $N$ is the number of individuals in the training set, $N_{\text{test}}$ is the number of individuals in the test set, $N_{\text{obs}}$ is the number of observations in the training set, $M$ is the number of SNPs used, $\hat{r}$ is the estimated GP accuracy, $\hat{\lambda}$ is the scaling factor used in finding $\hat{r}$ as described in the main text, $\overline{G}_{\text{ac}}$ is the mean relatedness between the training and test set individuals $\tilde{G}_{\text{ac}}$ is the precision of the relatedness between training and test set individuals and $d$ is the value that was added to the diagonal of the GRM for the respective model.

| Scenario | $N$ | $N_{\text{test}}$ | $N_{\text{obs}}$ | $M$ | $\hat{r}$ | $\hat{\lambda}$ | $\overline{G}_{\text{ac}}$ | $\tilde{G}_{\text{ac}}$ | $d$ |
|---|---|---|---|---|---|---|---|---|---|
| Helgeland | 3081 | 343 | 7179 | 65220 | 0.56 | 0.56 | -3.0E-04 | 1011.74 | 6.6E-03 |
| Helgeland | 3082 | 342 | 7150 | 65220 | 0.45 | 0.54 | -2.9E-04 | 986.15 | 6.6E-03 |
| Helgeland | 3082 | 342 | 7217 | 65220 | 0.74 | 0.52 | -3.2E-04 | 1028.35 | 6.6E-03 |
| Helgeland | 3081 | 343 | 7248 | 65220 | 0.93 | 0.51 | -3.2E-04 | 939.75 | 6.6E-03 |
| Helgeland | 3082 | 342 | 7222 | 65220 | 0.53 | 0.56 | -3.1E-04 | 981.62 | 6.6E-03 |
| Helgeland | 3082 | 342 | 7187 | 65220 | 0.52 | 0.53 | -3.0E-04 | 940.36 | 6.6E-03 |
| Helgeland | 3081 | 343 | 7202 | 65220 | 0.50 | 0.55 | -3.1E-04 | 991.61 | 6.6E-03 |
| Helgeland | 3082 | 342 | 7199 | 65220 | 0.62 | 0.58 | -2.9E-04 | 973.20 | 6.6E-03 |
| Helgeland | 3082 | 342 | 7198 | 65220 | 0.75 | 0.55 | -3.1E-04 | 978.97 | 6.6E-03 |
| Helgeland | 3081 | 343 | 7171 | 65220 | 0.98 | 0.47 | -3.1E-04 | 963.80 | 6.6E-03 |
| Southern | 2028 | 226 | 3755 | 64917 | 0.52 | 0.51 | -4.6E-04 | 959.22 | 5.0E-03 |
| Southern | 2029 | 225 | 3738 | 64917 | 0.57 | 0.52 | -5.0E-04 | 913.87 | 5.0E-03 |
| Southern | 2029 | 225 | 3753 | 64917 | 0.19 | 0.60 | -4.4E-04 | 936.63 | 5.0E-03 |
| Southern | 2028 | 226 | 3735 | 64917 | 0.59 | 0.54 | -4.5E-04 | 955.92 | 5.0E-03 |
| Southern | 2029 | 225 | 3752 | 64917 | 0.67 | 0.56 | -4.8E-04 | 955.69 | 5.0E-03 |
| Southern | 2029 | 225 | 3706 | 64917 | 0.60 | 0.56 | -4.7E-04 | 939.30 | 5.0E-03 |
| Southern | 2028 | 226 | 3764 | 64917 | 0.66 | 0.51 | -4.8E-04 | 952.82 | 5.0E-03 |
| Southern | 2029 | 225 | 3725 | 64917 | 0.86 | 0.49 | -4.7E-04 | 983.14 | 5.0E-03 |
| Southern | 2029 | 225 | 3686 | 64917 | 0.62 | 0.55 | -4.6E-04 | 882.99 | 5.0E-03 |
| Southern | 2028 | 226 | 3745 | 64917 | 0.72 | 0.49 | -4.5E-04 | 955.63 | 5.0E-03 |
| Merged | 5110 | 568 | 10874 | 64943 | 0.76 | 0.52 | -1.9E-04 | 1517.27 | 6.1E-03 |
| Merged | 5110 | 568 | 10942 | 64943 | 0.43 | 0.53 | -1.9E-04 | 1511.77 | 6.1E-03 |
| Merged | 5110 | 568 | 10893 | 64943 | 0.70 | 0.50 | -2.0E-04 | 1516.07 | 6.1E-03 |
| Merged | 5111 | 567 | 10941 | 64943 | 0.56 | 0.53 | -1.9E-04 | 1522.85 | 6.1E-03 |
| Merged | 5110 | 568 | 10938 | 64943 | 0.63 | 0.51 | -1.9E-04 | 1527.88 | 6.1E-03 |
| Merged | 5110 | 568 | 10992 | 64943 | 0.63 | 0.52 | -1.9E-04 | 1565.06 | 6.1E-03 |
| Merged | 5111 | 567 | 10929 | 64943 | 0.66 | 0.50 | -1.8E-04 | 1459.52 | 6.1E-03 |
| Merged | 5110 | 568 | 10951 | 64943 | 0.58 | 0.52 | -1.8E-04 | 1472.10 | 6.1E-03 |
| Merged | 5110 | 568 | 10969 | 64943 | 0.67 | 0.50 | -1.8E-04 | 1540.39 | 6.1E-03 |
| Merged | 5110 | 568 | 10903 | 64943 | 0.86 | 0.49 | -1.8E-04 | 1512.90 | 6.1E-03 |

#### Across-population GP: Body mass

Table S10: Statistics related to each across-population GP model with body mass as response. $N$ is the number of individuals in the training set, $N_{\text{test}}$ is the number of individuals in the test set, $N_{\text{obs}}$ is the number of observations in the training set, $M$ is the number of SNPs used, $\hat{r}$ is the estimated GP accuracy, $\hat{\lambda}$ is the scaling factor used in finding $\hat{r}$ as described in the main text, $\frac{h^2_{\text{GP}}}{h^2_{\text{AM}}}$ is the heritability ratio described in the main text, $\overline{G}_{\text{ac}}$ is the mean relatedness between the training and test set individuals $\tilde{G}_{\text{ac}}$ is the precision of the relatedness between training and test set individuals, $\Delta$LD1 is LD measure 1, $\Delta$LD2 is LD measure 2, $F_{st}$ is the population-pairwise fixation index between the training and test set, $\rho_g$ is the trait-specific estimated genetic correlation between the training and test set, $\rho_\varepsilon$ is the estimated residual (environmental) correlation between the training and test set and $d$ is the value that was added to the diagonal of the GRM for the respective model.

| Train islands | Test islands | $N$ | $N_{\text{test}}$ | $N_{\text{obs}}$ | $M$ | $\hat{r}$ | $\hat{\lambda}$ | $\frac{h^2_{\text{GP}}}{h^2_{\text{AM}}}$ | $\overline{G}_{\text{ac}}$ | $\tilde{G}_{\text{ac}}$ | $\Delta$LD1 | $\Delta$LD2 | $F_{st}$ | $\rho_g$ | $\rho_\varepsilon$ | $d$ |
|---|---|---|---|---|---|---|---|---|---|---|---|---|---|---|---|---|
| Helgeland | Nesøy | 3323 | 133 | 7544 | 65221 | 0.41 | 0.50 | 1.10 | -0.002 | 3173.80 | 0.0015 | 0.052 | 0.014 | 0.38 | 0.92 | 6.6E-03 |
| Helgeland | Myken | 3368 | 88 | 7837 | 65221 | 0.86 | 0.50 | 1.09 | -0.002 | 2278.34 | 0.0012 | 0.057 | 0.017 | 0.33 | 0.91 | 6.6E-03 |
| Helgeland | Træna | 3184 | 272 | 7616 | 65221 | 0.27 | 0.54 | 0.86 | -0.002 | 2307.47 | 0.0021 | 0.037 | 0.008 | 0.39 | 0.92 | 6.6E-03 |
| Helgeland | Selvær | 3234 | 222 | 7689 | 65221 | 0.32 | 0.55 | 0.91 | -0.002 | 2144.63 | 0.0017 | 0.039 | 0.008 | 0.45 | 0.92 | 6.6E-03 |
| Helgeland | Gjerøy | 2913 | 543 | 6534 | 65221 | 0.16 | 0.50 | 0.98 | -0.005 | 3550.86 | 0.0043 | 0.038 | 0.009 | 0.46 | 0.92 | 6.6E-03 |
| Helgeland | Hestmannøy | 2423 | 1033 | 5203 | 65221 | 0.38 | 0.50 | 1.03 | -0.005 | 3380.29 | 0.0045 | 0.030 | 0.006 | 0.31 | 0.92 | 6.6E-03 |
| Helgeland | Indre Kvarøy | 3085 | 371 | 7135 | 65221 | 0.49 | 0.49 | 1.02 | -0.004 | 2692.72 | 0.0031 | 0.040 | 0.010 | 0.47 | 0.92 | 6.6E-03 |
| Helgeland | Lurøy-Onøy | 3190 | 266 | 7362 | 65221 | 0.03 | 0.52 | 1.01 | -0.003 | 3068.63 | 0.0022 | 0.040 | 0.010 | 0.28 | 0.92 | 6.6E-03 |
| Helgeland | Lovund | 3309 | 147 | 7796 | 65221 | 0.55 | 0.52 | 0.94 | -0.001 | 2284.41 | 0.0010 | 0.036 | 0.006 | 0.36 | 0.92 | 6.6E-03 |
| Helgeland | Sleneset | 3267 | 189 | 7742 | 65221 | 0.37 | 0.57 | 0.78 | -0.003 | 2547.18 | 0.0021 | 0.046 | 0.012 | 0.55 | 0.91 | 6.6E-03 |
| Helgeland | Aldra | 3283 | 173 | 7354 | 65221 | 0.34 | 0.55 | 0.92 | -0.008 | 4908.19 | 0.0042 | 0.069 | 0.035 | -0.03 | 0.92 | 6.6E-03 |
| Helgeland | Non-farm | 2538 | 918 | 6772 | 65221 | 0.17 | 0.53 | 0.87 | -0.005 | 4315.31 | 0.0048 | 0.033 | 0.006 | 0.44 | 0.93 | 6.6E-03 |
| Helgeland | Farm | 937 | 2519 | 1247 | 65221 | 0.15 | 0.50 | 0.52 | -0.005 | 4196.60 | 0.0048 | 0.032 | 0.006 | 0.68 | 0.91 | 6.6E-03 |
| Non-farm | Nesøy | 918 | 133 | 1205 | 65221 | 0.48 | 0.46 | 0.83 | -0.001 | 3212.31 | 0.0054 | 0.055 | 0.016 | 0.62 | 0.91 | 1.0E-09 |
| Non-farm | Myken | 830 | 88 | 1065 | 65217 | 0.55 | 0.51 | 1.12 | 0.009 | 1122.54 | 0.0042 | 0.056 | 0.016 | 0.67 | 0.91 | 1.0E-09 |
| Non-farm | Træna | 646 | 272 | 844 | 65217 | 0.36 | 0.55 | 1.13 | 0.011 | 897.66 | 0.0049 | 0.034 | 0.005 | 0.65 | 0.92 | 1.0E-09 |
| Non-farm | Selvær | 696 | 222 | 917 | 65217 | 0.37 | 0.53 | 1.01 | 0.011 | 826.21 | 0.0046 | 0.035 | 0.006 | 0.71 | 0.92 | 1.0E-09 |
| Non-farm | Gjerøy | 918 | 543 | 1205 | 65220 | 0.06 | 0.51 | 0.50 | -0.008 | 6435.98 | 0.0099 | 0.045 | 0.013 | 0.64 | 0.93 | 1.0E-09 |
| Non-farm | Hestmannøy | 918 | 1033 | 1205 | 65221 | 0.15 | 0.50 | 0.50 | -0.006 | 4833.89 | 0.0076 | 0.038 | 0.009 | 0.66 | 0.92 | 1.0E-09 |
| Non-farm | Indre Kvarøy | 918 | 371 | 1205 | 65221 | 0.34 | 0.49 | 0.79 | -0.004 | 3531.31 | 0.0082 | 0.045 | 0.012 | 0.72 | 0.92 | 1.0E-09 |
| Non-farm | Lurøy-Onøy | 918 | 266 | 1205 | 65212 | -0.13 | 0.52 | 0.45 | -0.000 | 2837.14 | 0.0065 | 0.045 | 0.012 | 0.65 | 0.93 | 1.0E-09 |
| Non-farm | Lovund | 771 | 147 | 1024 | 65217 | 0.64 | 0.53 | 1.03 | 0.010 | 1129.47 | 0.0030 | 0.035 | 0.005 | 0.67 | 0.92 | 1.0E-09 |
| Non-farm | Sleneset | 729 | 189 | 970 | 65217 | 0.32 | 0.58 | 1.20 | 0.006 | 1483.43 | 0.0066 | 0.047 | 0.012 | 0.73 | 0.92 | 1.0E-09 |
| Non-farm | Aldra | 918 | 173 | 1205 | 65221 | 0.16 | 0.50 | 0.72 | -0.009 | 7107.91 | 0.0126 | 0.073 | 0.038 | 0.60 | 0.92 | 1.0E-09 |
| Non-farm | Farm | 918 | 2519 | 1205 | 65221 | 0.13 | 0.51 | 0.50 | -0.005 | 4341.38 | 0.0049 | 0.033 | 0.006 | 0.57 | 0.93 | 4.0E-03 |
| Farm | Nesøy | 2386 | 133 | 6297 | 65221 | 0.36 | 0.49 | 1.04 | -0.003 | 3169.44 | 0.0023 | 0.053 | 0.015 | 0.28 | 0.92 | 1.0E-09 |
| Farm | Myken | 2519 | 88 | 6730 | 65221 | 0.83 | 0.49 | 1.06 | -0.006 | 4193.67 | 0.0018 | 0.059 | 0.019 | 0.43 | 0.91 | 1.0E-09 |
| Farm | Træna | 2519 | 272 | 6730 | 65221 | 0.03 | 0.54 | 0.76 | -0.006 | 4927.24 | 0.0029 | 0.040 | 0.009 | 0.40 | 0.92 | 1.0E-09 |
| Farm | Selvær | 2519 | 222 | 6730 | 65221 | 0.00 | 0.54 | 0.83 | -0.006 | 4954.24 | 0.0026 | 0.042 | 0.010 | 0.37 | 0.92 | 1.0E-09 |
| Farm | Gjerøy | 1976 | 543 | 5287 | 65221 | 0.08 | 0.51 | 1.02 | -0.004 | 3047.01 | 0.0055 | 0.039 | 0.009 | 0.41 | 0.92 | 1.0E-09 |
| Farm | Hestmannøy | 1486 | 1033 | 3956 | 65221 | 0.34 | 0.50 | 0.91 | -0.004 | 2909.69 | 0.0056 | 0.032 | 0.006 | 0.46 | 0.92 | 1.0E-09 |
| Farm | Indre Kvarøy | 2148 | 371 | 5888 | 65221 | 0.39 | 0.50 | 0.81 | -0.004 | 2437.54 | 0.0043 | 0.041 | 0.010 | 0.54 | 0.92 | 1.0E-09 |
| Farm | Lurøy-Onøy | 2253 | 266 | 6115 | 65221 | 0.13 | 0.50 | 1.03 | -0.004 | 3202.95 | 0.0033 | 0.043 | 0.011 | 0.38 | 0.92 | 1.0E-09 |
| Farm | Lovund | 2519 | 147 | 6730 | 65221 | 0.13 | 0.53 | 0.70 | -0.004 | 3977.23 | 0.0016 | 0.040 | 0.008 | 0.36 | 0.92 | 1.0E-09 |
| Farm | Sleneset | 2519 | 189 | 6730 | 65221 | 0.25 | 0.59 | 0.91 | -0.006 | 3544.56 | 0.0029 | 0.048 | 0.014 | 0.16 | 0.92 | 1.0E-09 |
| Farm | Aldra | 2346 | 173 | 6107 | 65221 | 0.31 | 0.53 | 0.90 | -0.008 | 4381.58 | 0.0057 | 0.070 | 0.035 | 0.55 | 0.92 | 1.0E-09 |
| Farm | Non-farm | 2519 | 918 | 6730 | 65221 | 0.17 | 0.52 | 0.97 | -0.005 | 4341.38 | 0.0049 | 0.033 | 0.006 | 0.45 | 0.91 | 4.0E-03 |
| Southern | Leka | 1666 | 588 | 2947 | 64917 | 0.15 | 0.64 | 1.03 | -0.009 | 5010.31 | 0.0066 | 0.041 | 0.010 | 0.65 | 0.93 | 5.0E-03 |
| Southern | Vega | 1845 | 409 | 3259 | 64917 | 0.17 | 0.60 | 1.02 | -0.011 | 5598.08 | 0.0065 | 0.047 | 0.016 | 0.65 | 0.93 | 5.0E-03 |
| Southern | Vikna | 1668 | 586 | 3143 | 64917 | 0.33 | 0.61 | 1.09 | -0.006 | 6151.58 | 0.0048 | 0.035 | 0.007 | 0.58 | 0.94 | 5.0E-03 |
| Southern | Lauvøya | 1589 | 665 | 3036 | 64917 | 0.21 | 0.67 | 0.26 | -0.009 | 6462.24 | 0.0064 | 0.038 | 0.010 | 0.54 | 0.92 | 5.0E-03 |
| Helgeland | Southern | 3461 | 2249 | 7987 | 64943 | 0.10 | 0.60 | 0.73 | -0.009 | 12959.11 | 0.0066 | 0.034 | 0.009 | 0.36 | 0.93 | 6.2E-03 |
| Southern | Helgeland | 2268 | 3442 | 4156 | 64943 | 0.09 | 0.50 | 1.49 | -0.009 | 12168.65 | 0.0066 | 0.034 | 0.009 | 0.52 | 0.92 | 6.2E-03 |

#### Across-population GP: Tarsus length

Table S11: Statistics related to each across-population GP model with tarsus length as response. $N$ is the number of individuals in the training set, $N_{\text{test}}$ is the number of individuals in the test set, $N_{\text{obs}}$ is the number of observations in the training set, $M$ is the number of SNPs used, $\hat{r}$ is the estimated GP accuracy, $\hat{\lambda}$ is the scaling factor used in finding $\hat{r}$ as described in the main text, $\frac{h^2_{\text{GP}}}{h^2_{\text{AM}}}$ is the heritability ratio described in the main text, $\overline{G}_{\text{ac}}$ is the mean relatedness between the training and test set individuals $\tilde{G}_{\text{ac}}$ is the precision of the relatedness between training and test set individuals, $\Delta$LD1 is LD measure 1, $\Delta$LD2 is LD measure 2, $F_{st}$ is the population-pairwise fixation index between the training and test set, $\rho_g$ is the trait-specific estimated genetic correlation between the training and test set, $\rho_\varepsilon$ is the estimated residual (environmental) correlation between the training and test set and $d$ is the value that was added to the diagonal of the GRM for the respective model.

| Train islands | Test islands | $N$ | $N_{\text{test}}$ | $N_{\text{obs}}$ | $M$ | $\hat{r}$ | $\hat{\lambda}$ | $\frac{h^2_{\text{GP}}}{h^2_{\text{AM}}}$ | $\overline{G}_{\text{ac}}$ | $\tilde{G}_{\text{ac}}$ | $\Delta$LD1 | $\Delta$LD2 | $F_{st}$ | $\rho_g$ | $\rho_\varepsilon$ | $d$ |
|---|---|---|---|---|---|---|---|---|---|---|---|---|---|---|---|---|
| Helgeland | Nesøy | 3314 | 131 | 7640 | 65221 | 0.15 | 0.75 | 0.86 | -0.002 | 3148.54 | 0.0015 | 0.052 | 0.014 | 0.44 | 0.93 | 6.6E-03 |
| Helgeland | Myken | 3356 | 89 | 7927 | 65221 | 0.36 | 0.70 | 0.82 | -0.002 | 2239.83 | 0.0012 | 0.057 | 0.017 | 0.51 | 0.96 | 6.6E-03 |
| Helgeland | Træna | 3176 | 269 | 7698 | 65221 | 0.32 | 0.71 | 0.75 | -0.002 | 2295.74 | 0.0020 | 0.037 | 0.008 | 0.38 | 0.91 | 6.6E-03 |
| Helgeland | Selvær | 3223 | 222 | 7772 | 65221 | 0.27 | 0.74 | 0.76 | -0.002 | 2108.90 | 0.0016 | 0.038 | 0.008 | 0.27 | 0.91 | 6.6E-03 |
| Helgeland | Gjerøy | 2901 | 544 | 6607 | 65221 | 0.40 | 0.55 | 1.51 | -0.005 | 3568.50 | 0.0043 | 0.038 | 0.009 | 0.48 | 0.89 | 6.6E-03 |
| Helgeland | Hestmannøy | 2423 | 1022 | 5264 | 65221 | 0.31 | 0.66 | 0.51 | -0.005 | 3385.84 | 0.0045 | 0.030 | 0.006 | 0.40 | 0.92 | 6.6E-03 |
| Helgeland | Indre Kvarøy | 3075 | 370 | 7188 | 65221 | 0.49 | 0.65 | 0.92 | -0.004 | 2696.11 | 0.0031 | 0.040 | 0.009 | 0.50 | 0.92 | 6.6E-03 |
| Helgeland | Lurøy-Onøy | 3178 | 267 | 7467 | 65221 | 0.26 | 0.59 | 1.20 | -0.003 | 3030.53 | 0.0022 | 0.040 | 0.010 | 0.43 | 0.94 | 6.6E-03 |
| Helgeland | Lovund | 3297 | 148 | 7882 | 65221 | 0.58 | 0.65 | 0.91 | -0.001 | 2267.87 | 0.0009 | 0.037 | 0.007 | 0.22 | 0.94 | 6.6E-03 |
| Helgeland | Sleneset | 3256 | 189 | 7820 | 65221 | 0.50 | 0.68 | 0.81 | -0.003 | 2550.02 | 0.0021 | 0.046 | 0.012 | 0.25 | 0.94 | 6.6E-03 |
| Helgeland | Aldra | 3270 | 175 | 7449 | 65221 | 0.36 | 0.69 | 0.92 | -0.009 | 4911.51 | 0.0043 | 0.069 | 0.035 | 0.39 | 0.93 | 6.6E-03 |
| Helgeland | Non-farm | 2528 | 917 | 6831 | 65221 | 0.25 | 0.70 | 0.78 | -0.005 | 4285.73 | 0.0048 | 0.033 | 0.006 | 0.47 | 0.91 | 6.6E-03 |
| Helgeland | Farm | 936 | 2509 | 1280 | 65221 | 0.14 | 0.62 | 0.15 | -0.005 | 4168.77 | 0.0047 | 0.032 | 0.006 | 0.58 | 0.90 | 6.6E-03 |
| Non-farm | Nesøy | 917 | 131 | 1236 | 65221 | 0.18 | 0.71 | 0.39 | -0.000 | 2973.36 | 0.0055 | 0.055 | 0.016 | 0.71 | 0.91 | 1.0E-09 |
| Non-farm | Myken | 828 | 89 | 1096 | 65217 | 0.38 | 0.73 | 1.54 | 0.009 | 1088.02 | 0.0044 | 0.056 | 0.016 | 0.63 | 0.96 | 1.0E-09 |
| Non-farm | Træna | 648 | 269 | 867 | 65217 | 0.25 | 0.76 | 1.45 | 0.011 | 892.76 | 0.0047 | 0.034 | 0.005 | 0.63 | 0.89 | 1.0E-09 |
| Non-farm | Selvær | 695 | 222 | 941 | 65217 | 0.27 | 0.75 | 1.41 | 0.011 | 817.33 | 0.0044 | 0.035 | 0.005 | 0.53 | 0.91 | 1.0E-09 |
| Non-farm | Gjerøy | 917 | 544 | 1236 | 65220 | 0.02 | 0.56 | 0.24 | -0.008 | 6407.81 | 0.0100 | 0.045 | 0.013 | 0.59 | 0.92 | 1.0E-09 |
| Non-farm | Hestmannøy | 917 | 1022 | 1236 | 65221 | 0.08 | 0.64 | 0.13 | -0.006 | 4825.20 | 0.0076 | 0.038 | 0.009 | 0.64 | 0.91 | 1.0E-09 |
| Non-farm | Indre Kvarøy | 917 | 370 | 1236 | 65221 | 0.32 | 0.67 | 0.35 | -0.004 | 3559.23 | 0.0082 | 0.045 | 0.012 | 0.67 | 0.93 | 1.0E-09 |
| Non-farm | Lurøy-Onøy | 917 | 267 | 1236 | 65212 | 0.01 | 0.57 | 0.21 | -0.000 | 2819.28 | 0.0067 | 0.045 | 0.012 | 0.71 | 0.94 | 1.0E-09 |
| Non-farm | Lovund | 769 | 148 | 1051 | 65217 | 0.47 | 0.68 | 0.69 | 0.010 | 1116.33 | 0.0031 | 0.035 | 0.005 | 0.60 | 0.94 | 1.0E-09 |
| Non-farm | Sleneset | 728 | 189 | 989 | 65217 | 0.38 | 0.69 | 1.63 | 0.006 | 1479.48 | 0.0066 | 0.047 | 0.012 | 0.46 | 0.94 | 1.0E-09 |
| Non-farm | Aldra | 917 | 175 | 1236 | 65221 | 0.21 | 0.66 | 0.25 | -0.010 | 7105.12 | 0.0128 | 0.073 | 0.038 | 0.71 | 0.94 | 1.0E-09 |
| Non-farm | Farm | 917 | 2509 | 1236 | 65221 | 0.10 | 0.62 | 0.15 | -0.005 | 4311.29 | 0.0048 | 0.033 | 0.006 | 0.53 | 0.91 | 4.0E-03 |
| Farm | Nesøy | 2378 | 131 | 6360 | 65221 | 0.12 | 0.72 | 0.90 | -0.003 | 3233.92 | 0.0023 | 0.053 | 0.015 | 0.52 | 0.92 | 1.0E-09 |
| Farm | Myken | 2509 | 89 | 6787 | 65221 | 0.24 | 0.70 | 1.13 | -0.006 | 4203.09 | 0.0019 | 0.059 | 0.019 | 0.43 | 0.94 | 1.0E-09 |
| Farm | Træna | 2509 | 269 | 6787 | 65221 | 0.24 | 0.73 | 1.10 | -0.006 | 4925.65 | 0.0029 | 0.041 | 0.010 | 0.54 | 0.92 | 1.0E-09 |
| Farm | Selvær | 2509 | 222 | 6787 | 65221 | 0.07 | 0.74 | 0.93 | -0.006 | 4803.36 | 0.0025 | 0.042 | 0.010 | 0.53 | 0.89 | 1.0E-09 |
| Farm | Gjerøy | 1965 | 544 | 5327 | 65221 | 0.36 | 0.57 | 1.15 | -0.004 | 3066.64 | 0.0055 | 0.039 | 0.009 | 0.25 | 0.93 | 1.0E-09 |
| Farm | Hestmannøy | 1487 | 1022 | 3984 | 65221 | 0.31 | 0.64 | 0.53 | -0.004 | 2919.50 | 0.0057 | 0.032 | 0.006 | 0.52 | 0.91 | 1.0E-09 |
| Farm | Indre Kvarøy | 2139 | 370 | 5908 | 65221 | 0.39 | 0.67 | 0.84 | -0.004 | 2435.04 | 0.0042 | 0.041 | 0.010 | 0.47 | 0.93 | 1.0E-09 |
| Farm | Lurøy-Onøy | 2242 | 267 | 6187 | 65221 | 0.37 | 0.62 | 1.00 | -0.004 | 3153.37 | 0.0033 | 0.043 | 0.011 | 0.56 | 0.95 | 1.0E-09 |
| Farm | Lovund | 2509 | 148 | 6787 | 65221 | 0.41 | 0.69 | 0.84 | -0.005 | 3968.44 | 0.0015 | 0.040 | 0.008 | 0.44 | 0.96 | 1.0E-09 |
| Farm | Sleneset | 2509 | 189 | 6787 | 65221 | 0.35 | 0.73 | 1.01 | -0.005 | 3550.59 | 0.0030 | 0.048 | 0.014 | 0.54 | 0.90 | 1.0E-09 |
| Farm | Aldra | 2334 | 175 | 6169 | 65221 | 0.34 | 0.69 | 0.96 | -0.008 | 4383.92 | 0.0058 | 0.070 | 0.036 | 0.42 | 0.95 | 1.0E-09 |
| Farm | Non-farm | 2509 | 917 | 6787 | 65221 | 0.24 | 0.72 | 1.04 | -0.005 | 4311.29 | 0.0048 | 0.033 | 0.006 | 0.38 | 0.91 | 4.0E-03 |
| Southern | Leka | 1667 | 588 | 2993 | 64917 | 0.15 | 0.70 | 0.86 | -0.009 | 5008.84 | 0.0066 | 0.041 | 0.010 | 0.54 | 0.91 | 5.0E-03 |
| Southern | Vega | 1847 | 408 | 3344 | 64917 | 0.12 | 0.67 | 0.98 | -0.011 | 5599.24 | 0.0065 | 0.047 | 0.016 | 0.51 | 0.91 | 5.0E-03 |
| Southern | Vikna | 1667 | 588 | 3182 | 64917 | 0.16 | 0.64 | 1.04 | -0.006 | 6154.38 | 0.0048 | 0.035 | 0.007 | 0.53 | 0.91 | 5.0E-03 |
| Southern | Lauvøya | 1590 | 665 | 3007 | 64917 | 0.15 | 0.73 | 0.13 | -0.009 | 6468.81 | 0.0063 | 0.038 | 0.010 | 0.51 | 0.87 | 5.0E-03 |
| Helgeland | Southern | 3450 | 2250 | 8077 | 64943 | 0.15 | 0.63 | 0.99 | -0.009 | 12956.45 | 0.0066 | 0.034 | 0.009 | 0.22 | 0.94 | 6.1E-03 |
| Southern | Helgeland | 2269 | 3431 | 4205 | 64943 | 0.07 | 0.65 | 1.00 | -0.009 | 12165.15 | 0.0066 | 0.034 | 0.009 | 0.37 | 0.89 | 6.1E-03 |

### Across-population GP: Wing length

Table S12: Statistics related to each across-population GP model with wing length as response. $N$ is the number of individuals in the training set, $N_{\text{test}}$ is the number of individuals in the test set, $N_{\text{obs}}$ is the number of observations in the training set, $M$ is the number of SNPs used, $\hat{r}$ is the estimated GP accuracy, $\hat{\lambda}$ is the scaling factor used in finding $\hat{r}$ as described in the main text, $\frac{h^2_{\text{GP}}}{h^2_{\text{AM}}}$ is the heritability ratio described in the main text, $\overline{G}_{\text{ac}}$ is the mean relatedness between the training and test set individuals $\tilde{G}_{\text{ac}}$ is the precision of the relatedness between training and test set individuals, $\Delta$LD1 is LD measure 1, $\Delta$LD2 is LD measure 2, $F_{st}$ is the population-pairwise fixation index between the training and test set, $\rho_g$ is the trait-specific estimated genetic correlation between the training and test set, $\rho_\varepsilon$ is the estimated residual (environmental) correlation between the training and test set and $d$ is the value that was added to the diagonal of the GRM for the respective model.

| Train islands | Test islands | $N$ | $N_{\text{test}}$ | $N_{\text{obs}}$ | $M$ | $\hat{r}$ | $\hat{\lambda}$ | $\frac{h^2_{\text{GP}}}{h^2_{\text{AM}}}$ | $\overline{G}_{\text{ac}}$ | $\tilde{G}_{\text{ac}}$ | $\Delta$LD1 | $\Delta$LD2 | $F_{st}$ | $\rho_g$ | $\rho_\varepsilon$ | $d$ |
|---|---|---|---|---|---|---|---|---|---|---|---|---|---|---|---|---|
| Helgeland | Nesøy | 3292 | 132 | 7569 | 65220 | 0.16 | 0.56 | 0.93 | -0.002 | 3154.59 | 0.0015 | 0.052 | 0.014 | 0.15 | 0.91 | 6.6E-03 |
| Helgeland | Myken | 3335 | 89 | 7857 | 65220 | 0.81 | 0.47 | 1.25 | -0.002 | 2233.41 | 0.0012 | 0.057 | 0.017 | 0.27 | 0.91 | 6.6E-03 |
| Helgeland | Træna | 3157 | 267 | 7634 | 65220 | 0.26 | 0.55 | 1.02 | -0.002 | 2316.74 | 0.0020 | 0.037 | 0.008 | 0.17 | 0.91 | 6.6E-03 |
| Helgeland | Selvær | 3204 | 220 | 7705 | 65220 | 0.62 | 0.50 | 1.06 | -0.002 | 2123.52 | 0.0016 | 0.038 | 0.008 | 0.23 | 0.92 | 6.6E-03 |
| Helgeland | Gjerøy | 2883 | 541 | 6547 | 65220 | 0.26 | 0.56 | 0.97 | -0.005 | 3573.12 | 0.0043 | 0.038 | 0.009 | 0.26 | 0.92 | 6.6E-03 |
| Helgeland | Hestmannøy | 2409 | 1015 | 5232 | 65220 | 0.27 | 0.56 | 0.77 | -0.005 | 3380.30 | 0.0046 | 0.030 | 0.006 | 0.17 | 0.90 | 6.6E-03 |
| Helgeland | Indre Kvarøy | 3057 | 367 | 7119 | 65220 | 0.57 | 0.52 | 0.99 | -0.004 | 2701.24 | 0.0031 | 0.040 | 0.009 | 0.18 | 0.92 | 6.6E-03 |
| Helgeland | Lurøy-Onøy | 3158 | 266 | 7414 | 65220 | 0.68 | 0.54 | 0.91 | -0.003 | 3021.66 | 0.0021 | 0.040 | 0.010 | 0.22 | 0.91 | 6.6E-03 |
| Helgeland | Lovund | 3276 | 148 | 7813 | 65220 | 0.40 | 0.54 | 1.11 | -0.001 | 2274.08 | 0.0010 | 0.037 | 0.007 | 0.11 | 0.91 | 6.6E-03 |
| Helgeland | Sleneset | 3238 | 186 | 7752 | 65220 | 0.36 | 0.55 | 0.90 | -0.003 | 2547.44 | 0.0022 | 0.046 | 0.012 | 0.34 | 0.91 | 6.6E-03 |
| Helgeland | Aldra | 3250 | 174 | 7372 | 65220 | 0.21 | 0.51 | 1.09 | -0.008 | 4900.34 | 0.0043 | 0.069 | 0.035 | 0.33 | 0.91 | 6.6E-03 |
| Helgeland | Non-farm | 2514 | 910 | 6773 | 65220 | 0.27 | 0.53 | 1.06 | -0.005 | 4274.74 | 0.0048 | 0.033 | 0.006 | 0.16 | 0.92 | 6.6E-03 |
| Helgeland | Farm | 929 | 2495 | 1268 | 65220 | 0.10 | 0.54 | 0.45 | -0.005 | 4157.16 | 0.0047 | 0.032 | 0.006 | 0.49 | 0.89 | 6.6E-03 |
| Non-farm | Nesøy | 910 | 132 | 1224 | 65220 | 0.11 | 0.57 | 0.57 | -0.000 | 2971.26 | 0.0054 | 0.055 | 0.016 | 0.52 | 0.91 | 1.0E-09 |
| Non-farm | Myken | 821 | 89 | 1084 | 65216 | 0.43 | 0.50 | 1.09 | 0.009 | 1082.63 | 0.0044 | 0.056 | 0.016 | 0.51 | 0.92 | 1.0E-09 |
| Non-farm | Træna | 643 | 267 | 861 | 65216 | 0.28 | 0.55 | 1.03 | 0.011 | 907.01 | 0.0048 | 0.034 | 0.006 | 0.50 | 0.90 | 1.0E-09 |
| Non-farm | Selvær | 690 | 220 | 932 | 65216 | 0.59 | 0.52 | 0.84 | 0.011 | 828.17 | 0.0044 | 0.035 | 0.006 | 0.56 | 0.92 | 1.0E-09 |
| Non-farm | Gjerøy | 910 | 541 | 1224 | 65219 | -0.07 | 0.56 | 0.48 | -0.008 | 6390.60 | 0.0100 | 0.045 | 0.013 | 0.62 | 0.91 | 1.0E-09 |
| Non-farm | Hestmannøy | 910 | 1015 | 1224 | 65220 | 0.05 | 0.52 | 0.44 | -0.006 | 4817.53 | 0.0076 | 0.038 | 0.009 | 0.54 | 0.91 | 1.0E-09 |
| Non-farm | Indre Kvarøy | 910 | 367 | 1224 | 65220 | 0.10 | 0.52 | 0.83 | -0.004 | 3535.06 | 0.0082 | 0.045 | 0.012 | 0.68 | 0.91 | 1.0E-09 |
| Non-farm | Lurøy-Onøy | 910 | 266 | 1224 | 65212 | 0.53 | 0.54 | 0.42 | -0.000 | 2819.93 | 0.0067 | 0.045 | 0.012 | 0.32 | 0.91 | 1.0E-09 |
| Non-farm | Lovund | 762 | 148 | 1040 | 65216 | 0.34 | 0.51 | 0.90 | 0.010 | 1121.73 | 0.0032 | 0.036 | 0.005 | 0.52 | 0.92 | 1.0E-09 |
| Non-farm | Sleneset | 724 | 186 | 979 | 65216 | 0.34 | 0.60 | 1.23 | 0.006 | 1479.89 | 0.0066 | 0.047 | 0.012 | 0.46 | 0.92 | 1.0E-09 |
| Non-farm | Aldra | 910 | 174 | 1224 | 65220 | 0.12 | 0.55 | 0.54 | -0.010 | 7095.88 | 0.0130 | 0.073 | 0.038 | 0.53 | 0.92 | 1.0E-09 |
| Non-farm | Farm | 910 | 2495 | 1224 | 65220 | 0.06 | 0.55 | 0.43 | -0.005 | 4300.35 | 0.0048 | 0.033 | 0.006 | 0.60 | 0.90 | 3.9E-03 |
| Farm | Nesøy | 2363 | 132 | 6301 | 65220 | 0.16 | 0.54 | 1.12 | -0.003 | 3243.48 | 0.0023 | 0.053 | 0.015 | 0.39 | 0.91 | 1.0E-09 |
| Farm | Myken | 2495 | 89 | 6729 | 65220 | 0.68 | 0.49 | 1.18 | -0.006 | 4195.79 | 0.0018 | 0.059 | 0.019 | 0.27 | 0.92 | 1.0E-09 |
| Farm | Træna | 2495 | 267 | 6729 | 65220 | 0.09 | 0.54 | 0.79 | -0.006 | 4915.69 | 0.0029 | 0.041 | 0.009 | 0.30 | 0.91 | 1.0E-09 |
| Farm | Selvær | 2495 | 220 | 6729 | 65220 | 0.32 | 0.52 | 0.90 | -0.006 | 4777.19 | 0.0025 | 0.042 | 0.010 | 0.14 | 0.91 | 1.0E-09 |
| Farm | Gjerøy | 1954 | 541 | 5279 | 65220 | 0.29 | 0.54 | 1.16 | -0.004 | 3074.25 | 0.0055 | 0.039 | 0.009 | 0.50 | 0.92 | 1.0E-09 |
| Farm | Hestmannøy | 1480 | 1015 | 3964 | 65220 | 0.29 | 0.55 | 0.79 | -0.004 | 2916.43 | 0.0057 | 0.032 | 0.006 | 0.53 | 0.91 | 1.0E-09 |
| Farm | Indre Kvarøy | 2128 | 367 | 5851 | 65220 | 0.58 | 0.52 | 1.07 | -0.004 | 2446.65 | 0.0043 | 0.041 | 0.010 | 0.43 | 0.92 | 1.0E-09 |
| Farm | Lurøy-Onøy | 2229 | 266 | 6146 | 65220 | 0.57 | 0.56 | 1.02 | -0.004 | 3139.56 | 0.0033 | 0.043 | 0.011 | 0.40 | 0.91 | 1.0E-09 |
| Farm | Lovund | 2495 | 148 | 6729 | 65220 | 0.31 | 0.55 | 0.85 | -0.005 | 3959.41 | 0.0015 | 0.040 | 0.008 | 0.30 | 0.92 | 1.0E-09 |
| Farm | Sleneset | 2495 | 186 | 6729 | 65220 | 0.26 | 0.55 | 1.32 | -0.005 | 3543.74 | 0.0029 | 0.048 | 0.014 | 0.28 | 0.92 | 1.0E-09 |
| Farm | Aldra | 2321 | 174 | 6104 | 65220 | 0.18 | 0.49 | 1.12 | -0.008 | 4373.07 | 0.0058 | 0.070 | 0.036 | 0.45 | 0.91 | 1.0E-09 |
| Farm | Non-farm | 2495 | 910 | 6729 | 65220 | 0.28 | 0.52 | 1.10 | -0.005 | 4300.35 | 0.0048 | 0.033 | 0.006 | 0.46 | 0.91 | 3.9E-03 |
| Southern | Leka | 1668 | 586 | 2977 | 64917 | 0.25 | 0.56 | 0.94 | -0.009 | 5007.22 | 0.0066 | 0.041 | 0.010 | 0.47 | 0.92 | 5.0E-03 |
| Southern | Vega | 1845 | 409 | 3324 | 64917 | 0.06 | 0.53 | 0.93 | -0.011 | 5602.19 | 0.0065 | 0.047 | 0.016 | 0.50 | 0.91 | 5.0E-03 |
| Southern | Vikna | 1666 | 588 | 3161 | 64917 | 0.12 | 0.50 | 1.11 | -0.006 | 6151.02 | 0.0049 | 0.035 | 0.007 | 0.50 | 0.91 | 5.0E-03 |
| Southern | Lauvøya | 1589 | 665 | 3004 | 64917 | 0.09 | 0.54 | 0.30 | -0.009 | 6475.60 | 0.0064 | 0.038 | 0.010 | 0.59 | 0.91 | 5.0E-03 |
| Helgeland | Southern | 3429 | 2249 | 8007 | 64943 | 0.17 | 0.55 | 0.93 | -0.009 | 12958.62 | 0.0066 | 0.034 | 0.009 | 0.15 | 0.91 | 6.1E-03 |
| Southern | Helgeland | 2268 | 3410 | 4185 | 64943 | 0.14 | 0.50 | 0.97 | -0.009 | 12163.28 | 0.0066 | 0.034 | 0.009 | 0.32 | 0.89 | 6.1E-03 |
